# Diverse patterns of secondary structure across genes and transposable elements are associated with siRNA production and epigenetic fate

**DOI:** 10.1101/2022.10.17.512609

**Authors:** Galen Martin, Edwin Solares, Aline Muyle, Alexandros Bousios, Brandon S. Gaut

**Affiliations:** Department of Ecology and Evolutionary Biology, University of California, Irvine; Department of Ecology and Evolutionary Biology, University of California, Davis; CEFE, Univ Montpellier, CNRS, EPHE, IRD, Montpellier, France; School of Life Sciences, University of Sussex, Brighton, UK

**Keywords:** small RNA, Zea mays, RNA interference, secondary structure, RNA directed DNA methylation, transposable elements, epigenome

## Abstract

RNA molecules carry information in their primary sequence and also their secondary structure. Secondary structure can confer important functional information, but it is also a potential signal for an RNAi-like host epigenetic response mediated by small interfering RNAs (siRNAs). In this study, we predicted local secondary structures in features of the maize genome, focusing on small regions that had folding energies similar to pre-miRNA loci. We found secondary structures to be common in retrotransposons, in *Helitrons*, and in genes. These structured regions mapped higher diversities of siRNAs than regions without structure, explaining up to 24% of variation of the siRNA distribution across some TE types. Among genes, those with secondary structure were 1.5-fold more highly expressed, on average, than genes without secondary structure. However, these genes were also more variably expressed across the 26 NAM lines, and this variability correlated with the number of mapping siRNAs. We conclude that local stem-loop structures are a nearly ubiquitous feature of expressed regions of the maize genome, that they correlate with higher siRNA mapping, and that they can represent a trade-off between functional need and the potentially negative consequences of siRNA production.

## INTRODUCTION

In a highly simplified view, plant genomes are composed of transposable elements (TEs) and genes. Both of these components use RNA to transmit coding information between one state (DNA) to another (protein). These RNA molecules carry information in their primary sequence of bases but also by their shape, or secondary structure. This shape is defined by bonds between the RNA bases of a transcript, and it can play a major role in mediating the relationship between genotype and phenotype because it affects the localization (Bullock et al., 2010), splicing (Buratti & Baralle, 2004), and translation (Ding et al., 2014) of mRNAs. As a result, secondary structure influences nearly every processing step in the life cycle of transcripts (Vandivier et al., 2016).

Hairpin-like secondary structures can also cause transcripts to enter the RNA interference (*RNAi*) pathway (Baulcombe 2004; Li et al., 2012), which degrades RNA molecules into small (21–24-nucleotide) RNAs. Entrance into RNAi occurs through the binding of *Dicer-like* proteins (DCL) (Fukudome & Fukuhara 2017) which specifically degrade double-stranded RNA (dsRNA). When single-stranded RNA (ssRNA) forms a hairpin-like secondary structure, DCLs recognize structured ssRNA as dsRNA, leading to the production of small RNAs. This mechanism is essential for the biogenesis of microRNAs (miRNAs), a class of small RNAs that are generally ~22-nt in length and that are derived from pre-miRNA genes with strong hairpin secondary structures. Once generated, miRNAs bind and transiently repress target genes through RNAi. In effect, miRNAs prime themselves–i.e., they ‘self-prime’– for degradation into small RNAs through their secondary structure.

While miRNAs transiently downregulate genes, a separate class of small RNAs – termed small interfering RNAs (siRNAs) – typically silence TEs. These siRNAs are distinguished from miRNAs by often having perfect complementarity to target sequences, and they direct both post-transcriptional gene silencing (PTGS) and transcriptional gene silencing (TGS) of TEs. In the PTGS pathway, siRNAs derived from dsRNA or hairpin RNA bind complementary transcripts and act as a primer for *RNA-dependent RNA polymerase VI* (RDR6), which copies the ssRNA transcript into dsRNA (Marí-Ordóñez et al., 2013; Matzke and Mosher, 2014; Nuthikattu et al., 2013). Subsequently, DCLs cut dsRNA into more siRNAs, which guide *ARGONAUTE* endoribonuclease enzymes to cleave complementary mRNA transcripts, establishing a feed-forward loop. This PTGS pathway eventually leads to TGS (Nuthikattu et al., 2013). When RNAi degrades TE transcripts – or TE-derived sequences, as in the *Mu-killer* locus (Slotkin et al., 2003) – the resulting siRNAs direct methyltransferase enzymes to complementary DNA sequences, initiating RNA-directed DNA methylation (RdDM) and epigenetic TGS of those genomic loci. Methylated, constitutively silenced genomic regions are then transcribed by separate RNA polymerases, RNA polymerase IV and V (Pol IV/V), into more siRNAs that further reinforce and spread methylation (Cuerda-Gil and Slotkin, 2016).

TGS and PTGS usually involve separate classes of siRNAs distinguished by sequence length (Borges and Martienssen, 2015). Typically, 21 and 22-nt siRNAs form through mRNA transcription by Pol II, the enzyme responsible for mRNA expression, and represent the products of PTGS. They are therefore dependent on the expression of a gene or a TE. In contrast, 24-nt siRNAs are products of Pol IV/V transcription in heterochromatic DNA, which are transcribed directly from genomic loci that have already been epigenetically silenced. These siRNAs reinforce silencing by directing methylation to homologous loci through RdDM (Matzke and Mosher, 2014). As such, 24-nt siRNA reflect TGS and are typically more abundant than 21 and 22-nt siRNAs. Therefore, one expects TEs with heavy 24-nt siRNA accumulation to be quiescent and TEs with heavy 21–22nt siRNA accumulation to experience at least some level of Pol II activity. However, this distinction is somewhat porous given that Pol IV/V can produce 21–22 nt siRNAs in some situations (Fultz & Slotkin, 2017; Panda et al., 2020).

Despite its importance, little is known about how host genomes distinguish TEs from genes and target *de novo* silencing. Several studies suggest that hairpin RNA structures act as an immune signal for *de novo* silencing of TEs (Slotkin et al., 2003; Sijen and Plasterk, 2003; Bousios et al., 2016; Hung & Slotkin 2021). One such example is *Mu-killer,* a locus that generates small RNAs that limits the activity of *Mu* elements in maize (*Zea mays* ssp. *mays*) (Slotkin et al., 2003). *Mu*-killer consists of a truncated, duplicated, and inverted copy of *Mu* that, when expressed, creates a folded substrate for *DCL* enzymes and is cut into trans-acting siRNAs that target active *Mu* transcripts. In this respect, *Mu-killer* is similar to miRNA biogenesis (O’Brien et al., 2018). Another potential example comes from Sirevirus long terminal repeat (LTR) retrotransposons in maize (Bousios et al., 2016), which occupy 20% of the B73 genome (Bousios 2012). In this study, the authors mapped siRNAs to full-length Sirevirus copies, reasoning that loci important for recognition and silencing should be associated with a larger number of siRNAs than other regions of the elements. Indeed, an excess of siRNAs mapped to clusters of palindromic motifs that defined a region with strong predicted secondary structure (Bousios et al., 2016). These studies present evidence that self-priming via secondary structure may direct silencing of at least some TEs; in fact, siRNAs derived from hairpin structures are frequent enough that they have their own name: hairpin RNAs or hpRNAs (Axtell, 2013).

If RNA sequences self-prime for siRNA production via RNA secondary structure, two important questions must be addressed. First, how common is this process across the diversity of TE categories? Thus far, its importance has been implicated in a few individual families or genera, but comparisons of secondary structure across TEs have not been performed. Second, secondary structure is not unique to TEs and exists within genes as well. What – if anything – prevents self-priming and degradation in these genes? Li et al. (2012) documented a positive relationship between mRNA structure and sRNA abundance for *Arabidopsis thaliana* genes, suggesting that genes are susceptible to self-priming. However, these genes are still expressed. Some countermeasures may moderate the potential effects of sRNAs on genes, including hypothesized protection against RNAi caused by high GC content (Hung and Slotkin 2021) and active gene demethylation (Gong et al., 2002; Zhang et al., 2018; Muyle et al., 2021). To our knowledge, however, there has not yet been a genome-wide comparison of secondary structure between genes and TEs; it is unknown whether TEs typically possess stronger secondary structures than genes.

In this study, we focus on the maize genome and explore relationships among predicted secondary structures in genes and TEs with small RNA targeting, gene expression, and chromatin accessibility (**Fig S1**). There are generally two ways to catalog secondary structure. The first is experimental, using approaches like SHAPE-seq, which can be applied to small regions or the transcribed component of whole genomes (Ding et al., 2013). This empirical approach is, however, difficult to perform on large genomes with high repeat content, and it also requires that the sequences of interest are expressed, preventing investigation of most plant TEs. The second method, which we adopted here, relies on bioinformatic predictions of minimum free energy (MFE) based on genome sequence data. A complication to this approach is that MFE is influenced by sequence length and composition, making it challenging to compare the strength of secondary structures across different features. To circumvent this issue, we predicted MFE in overlapping sliding windows of defined length (110 bp). We modeled these windows on the RNAfold predictions of pre-miRNA loci in a previous maize study (Wang et al., 2009), which showed that pre-miRNA windows of ~110 bp typically have MFEs <-40 kcal/mol and that a minimum MFE threshold of −40 kcal/mol can determine the ability of a ~110 nt pre-miRNA locus to form a hairpin structure. By focusing here on regions of similar size to these pre-miRNA transcripts, and by employing their threshold cutoff of of −40 kcal/mol, we in effect use miRNA loci as a ‘positive control’ for single-stranded RNAs that are known to self-prime into hairpin substrates, allowing us to test the quality of our computational inferences.

After performing computational annotation of the extent and folding strength of secondary structures in genes and TEs, we mapped siRNAs datasets from multiple libraries, allowing us to integrate secondary structure information with small RNA complement. With these data, we address three sets of questions. The first focuses on predicted secondary structure: How often do TEs and genes contain regions of strong predicted secondary structures? Are these secondary structures in specific locations of TEs and genes? And how do secondary structures compare between them? Our second set of questions focuses on the relationship between secondary structure and epigenetic features. Do regions of strong secondary structure correlate with siRNAs? If so, does this epigenetic feature differ between TEs and genes, and does this distinguish the active epigenetic state of genes from the silenced state of TEs? Finally, we synthesize genomic, transcription, and epigenetic data from 26 diverse maize lines to form a framework for interpreting the fate of genes, TEs, and their transcripts through the lens of secondary structure.

## RESULTS

### Measuring secondary structure in genomic features

We inferred local secondary structure across genome features of the B73 reference maize genome (version 4.0). The features included miRNA precursor loci, TEs and genes. The TEs included all families annotated by Jiao et al. (2017), and included both class I retroelements and class II DNA transposons. For both classes, we focused on superfamily categories (Wicker et al., 2007), which distinguished (for example) between *Ty3-Gypsy*/RLG and *Copia*/RLC LTR elements and among TIR elements like *Mutators*/DTM and *Harbingers*/DTH. [Note that throughout the paper we refer to TE superfamilies by their names and also their three-letter designation from Wicker et al., 2007 (**Table 1**).] For genes, we studied both the annotated gene – which included 5’ untranslated regions (UTRs), exons, introns and 3’ UTRs – as well as mature transcripts that lacked introns. Altogether, we examined 373,485 features representing 15 distinct categories (**Table 1**).

**Table 1.**
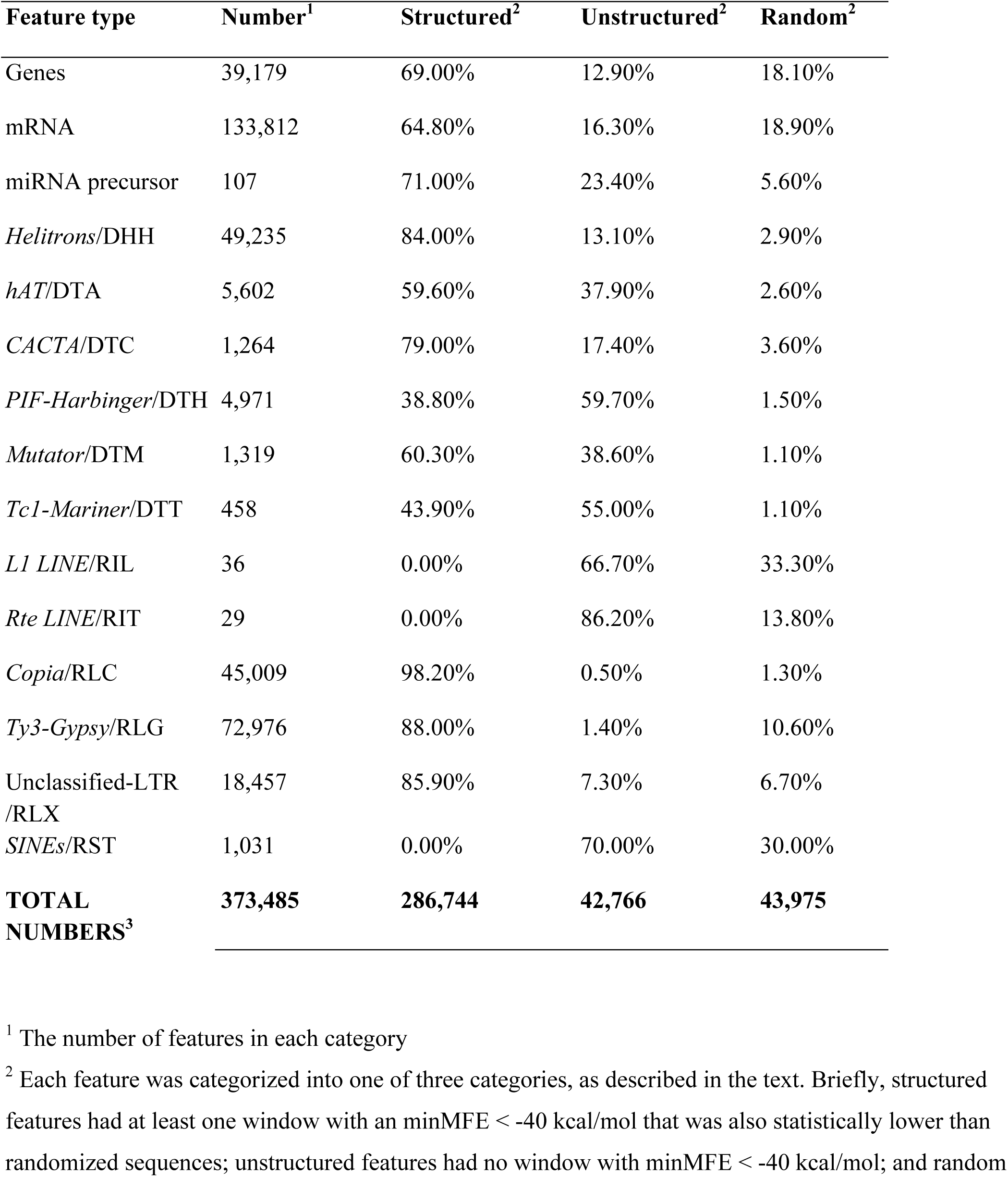
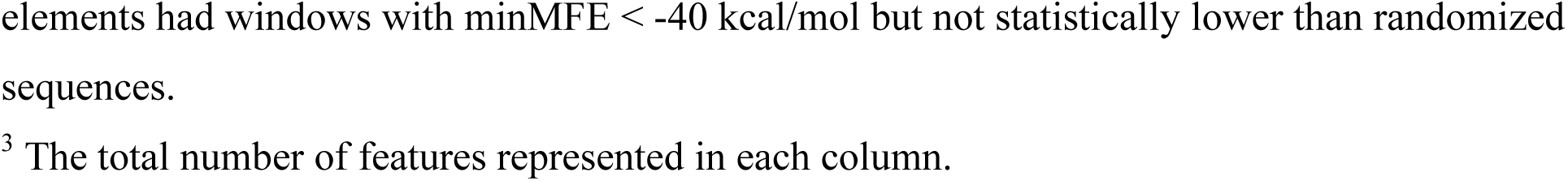
The percentage of each feature type determined to be structured, unstructured or random based on the sequence randomization and minMFE. See the primary text for definitions of the structure categories.

| Feature type | Number<sup>1</sup> | Structured<sup>2</sup> | Unstructured<sup>2</sup> | Random<sup>2</sup> |
| --- | --- | --- | --- | --- |
| Genes | 39,179 | 69.00% | 12.90% | 18.10% |
| mRNA | 133,812 | 64.80% | 16.30% | 18.90% |
| miRNA precursor | 107 | 71.00% | 23.40% | 5.60% |
| *Helitrons*/DHH | 49,235 | 84.00% | 13.10% | 2.90% |
| *hAT*/DTA | 5,602 | 59.60% | 37.90% | 2.60% |
| *CACTA*/DTC | 1,264 | 79.00% | 17.40% | 3.60% |
| *PIF-Harbinger*/DTH | 4,971 | 38.80% | 59.70% | 1.50% |
| *Mutator*/DTM | 1,319 | 60.30% | 38.60% | 1.10% |
| *Tc1-Mariner*/DTT | 458 | 43.90% | 55.00% | 1.10% |
| *L1 LINE*/RIL | 36 | 0.00% | 66.70% | 33.30% |
| *Rte LINE*/RIT | 29 | 0.00% | 86.20% | 13.80% |
| *Copia*/RLC | 45,009 | 98.20% | 0.50% | 1.30% |
| *Ty3-Gypsy*/RLG | 72,976 | 88.00% | 1.40% | 10.60% |
| Unclassified-LTR<br>/RLX | 18,457 | 85.90% | 7.30% | 6.70% |
| *SINEs*/RST | 1,031 | 0.00% | 70.00% | 30.00% |
| **TOTAL**<br>**NUMBERS<sup>3</sup>** | **373,485** | **286,744** | **42,766** | **43,975** |

<sup>1</sup> The number of features in each category
<sup>2</sup> Each feature was categorized into one of three categories, as described in the text. Briefly, structured features had at least one window with an minMFE < -40 kcal/mol that was also statistically lower than randomized sequences; unstructured features had no window with minMFE < -40 kcal/mol; and random
elements had windows with $\text{minMFE} < -40$ kcal/mol but not statistically lower than randomized sequences.
<sup>3</sup> The total number of features represented in each column.

For each feature, we predicted MFEs in sliding windows of 110 bp using RNAfold, mimicking previous work (Wang et al., 2009; Bousios et al., 2016). The advantages of this window-based approach were that we could use pre-miRNA transcript loci as positive controls, that we could explicitly compare results to the −40kcal/mol MFE threshold, that the output could be compared across features, and that it measured local folding properties across the length of specific features. Because each nucleotide of a feature corresponded to one sliding window (for all but the final 109 nucleotides of a sequence), predicting the MFE of windows was a massive bioinformatic undertaking. In total, we calculated the MFE for 3.56e9 windows.

Because each TE, gene or other feature consisted of many windows, we were able to characterize the secondary structure of each feature using multiple summary statistics. For example, we recorded the minimum MFE (minMFE), which we defined as the MFE of the window with the strongest predicted secondary structure for each feature (**Fig 1a**). We also calculated the mean MFE (meanMFE) across all windows of each feature (**Fig 1b**), the percentage of windows below the −40 kcal/mol threshold (propMFE) (**Fig 1c**), and the variance in MFE (varMFE) across all the windows within the feature (**Fig 1d**). minMFE pinpointed the MFE of the most structured window within a feature, while the other statistics summarized global MFE patterns across the feature.

**Figure 1:**
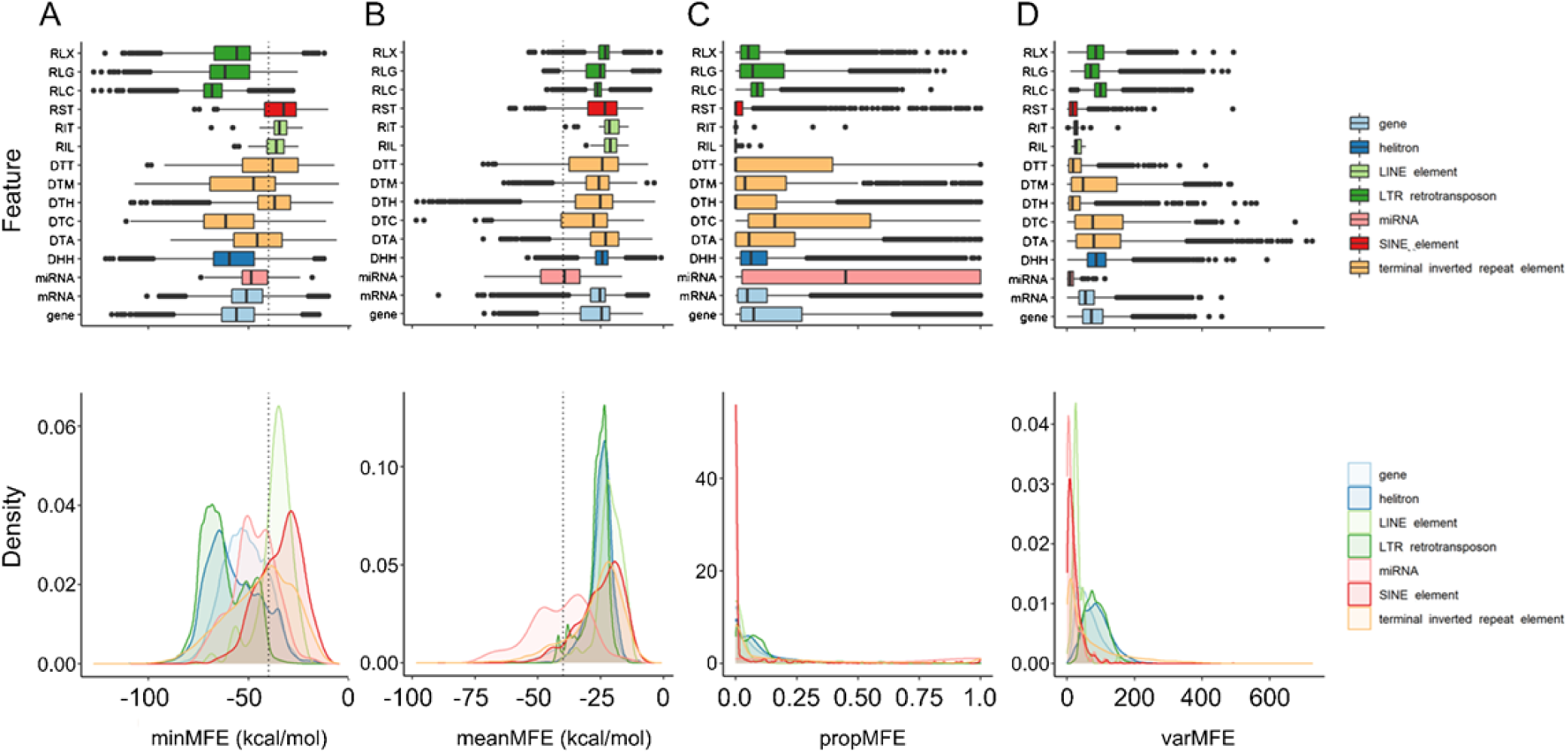
Variation in secondary structure between feature types. Each column represents a different statistic (see text) summarizing MFE in each feature type. The top row of box plots shows the statistic for each of the 15 feature groups defined in Table 1. The three letter codes represent types of TE superfamilies that are defined in Table 1. The bottom row plots the same information, but with the density for each of the groups defined by the color key. In both the box and density plots, the dotted line in minMFE and meanMFE delineates −40 kcal/mol, the cutoff point for windows with significant/miRNA-like secondary structure from Wang et al. (2009).

Two factors influence the stability of RNA secondary structure: base composition (e.g., higher GC content tends to induce more stable secondary structures) and primary sequence (e.g., whether the order of bases forms palindromes and stem-loop structures). Because we were primarily interested in secondary structure resulting from the latter, we controlled for base composition by randomizing the sequence of each feature five times and then repeated the MFE predictions each time, requiring another 5 x 3.56e9 window computations. By randomizing each feature, we identified features that had more stable secondary structures than expected given their nucleotide composition. We then classified a feature into one of three categories: i) “unstructured” when it had an minMFE > −40 kcal/mol, ii) “structured” when it had an minMFE < −40 kcal/mol and a significantly lower minMFE than the randomized permutations (p < 0.05, one-sided Wilcoxon test, Benjamini and Hochberg corrected) and iii) “random” when it had minMFE < −40 kcal/mol but did not have a significantly lower minMFE compared to permutations. We largely ignored the random category in downstream analyses, because it encompassed a group where strong secondary structure may be due to GC content rather than primary sequence. We report the differences between randomized and observed minMFE values for each feature category in **Fig S2**.

### Secondary structure is unevenly distributed among and within genomic features

Several broad patterns emerged from our summaries of secondary structure. First, in accordance with the results from Wang et al (2009), the majority of pre-miRNA loci (71.0%) were structured (**Figure 1a**; **Table 1**), suggesting that the −40k cal/mol cutoff is generally applicable for our secondary structure predictions. Second, by comparison, some categories had far lower percentages of structured features. For example, LINE/RIL&RIT and SINEs/RST, had no (0%) structured elements, and most (>70%) of these elements had no windows <-40k cal/mol (**Table 1**). Third, a few categories had higher percentages of structured features than miRNAs. For example, 98% of *Copia*/RLC elements were structured, which was not surprising given that most maize *Copia*/RLC elements are Sireviruses with known palindrome-derived secondary structures (Bousios et al., 2016) (**Table 1**). Additional categories with high percentages of structured features included *Ty3*/RLG elements (88%) and genes (69%). Note, however, that LTRs and genes also tended to be longer than the other features considered, and there was an overall negative relationship between feature length and minMFE (P < 2.2e-16, R2 = 0.20, linear model; **Fig S3**). Hence we did not completely circumvent the issue of length for some summary statistics, and so these comparisons of percentages need to be interpreted cautiously. Interestingly, however, the effect of length was weakest in genes, - i.e., both long and short genes tended to have similarly stable secondary structures (**Fig S3**).

Given variation in secondary structure among different categories of features, we next located regions of high secondary structure (i.e. regions below −40 kcal/mol) within each feature type. For these analyses, we focused only on the 286,836 structured features (**Table 1**) - i.e., the features with significant secondary structure as identified by sequence randomization. For each feature, we mapped the positions of low MFE regions along the length of the feature on a 0–1 scale from the 5’ start (0) to the 3’ end (1) (**Fig 2**). We first plotted these regions across TE superfamilies, beginning with the *Ty3-Gypsy*/RLG and *Copia*/RLC LTR retrotransposons that together constitute ~90% of TEs in the maize genome (Jiao et al., 2017). The MFE landscapes indicated strong signals of secondary structure within the LTRs region of *Copia*/RLC elements, consistent with the palindromic repeat regions of Sireviruses (Bousios et al., 2016). The low MFE regions of *Ty3-Gypsy*/RLG were in notably different locations compared to *Copia*/RLC; although there was some bias toward the 5’ region of TEs, there were no obvious peaks in *Ty3-Gypsy*/RLG LTRs. Overall, the *Ty3-Gypsy*/RLG LTRs had higher minMFEs (P < 0.01, t-test; **Fig 1**) than *Copia*/RLC with lower varMFE. For completeness, we also examined solo LTRs of LTR superfamilies, which mirrored the minMFE distributions within the LTRs of full length elements (**Fig S4**). In contrast to *Copia*/RLC elements, DNA transposon superfamilies had relatively uniform distributions of low MFE regions across their lengths, although the regions were biased slightly towards the edges of inverted repeats for TIR elements like *Mutator*/DTM (**Fig 2**), *hAT*/DTA and *CACTA*/DTC elements (**Fig. S3**). Lastly, *Helitrons*/DHH demonstrated a distinct bias towards their 3’ edge (**Fig 2**); 12% of annotated *Helitrons* had their lowest MFE region in the final 3’ window of the element, perhaps reflecting the ~11 nt stem-loop structure common to the *Helitron* 3’ end (Kapitonov & Jurka 2007; Xiong et al., 2014).

**Figure 2:**
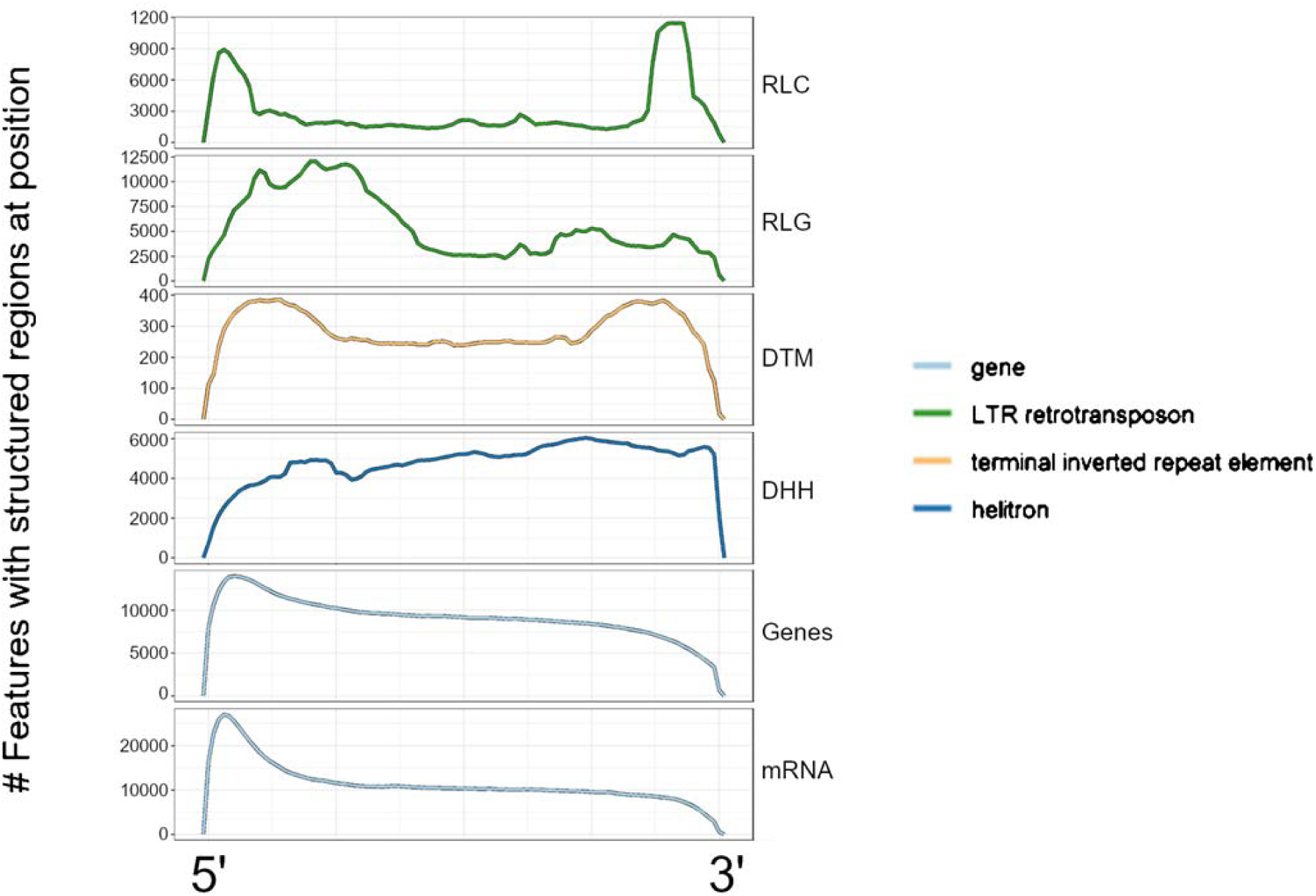
Landscapes of structured regions across feature types. Each row represents a metaprofile combining data from all members of each feature type. Features were divided into 100 equally sized bins from the 5’ end to the 3’ end, and the number of features with low MFE (<-40 kcal/mol) windows overlapping each of these bins was counted. A peak in the landscape represents a region of the feature type that often contains inferred stable secondary structure. All rows share the same x-axis, which is represented proportionally across the length of features, from 0.00 (5’ end) to 1.00 (3’ end).

These differences in the distribution of structure across the length of TEs could represent distinct structured sequence motifs. For each TE superfamily, we extracted all the sequences of structured regions (with all overlapping windows concatenated together) and input them into the Multiple EM for Motif Elicitation (MEME) suite motif discovery tool (Bailey and Elkan, 1994). As expected, we identified the previously identified consensus Sirevirus palindrome, CACCGGACtGTCCGGTG (**Fig S5**) as the most abundant motif in *Copia*/RLC elements (MEME e-value = 5.3e-677), appearing in 42.9% of RLC structured regions. Surprisingly, the same palindrome (**Fig S5**) was also the most abundant motif in *Helitron*/DHH transposons (MEME e-value = 1.0e-165), appearing in 5,231 DHH structured regions (10.7%). This observation could reflect a tendency for *Copia*/RLC elements to preferentially insert into *Helitron*/DHH elements, *Copia*/RLC capture and co-option by *Helitrons,* potential of TE borders, or even independent emergence of these motifs in the two superfamilies.

We also examined the distributions of secondary structure across genes and their mature transcripts/mRNAs (**Fig 2**). As expected (Li et al, 2012), secondary structure was biased towards the 5’ ends of genes and their mRNAs. In most genes, peaks of secondary structure were located within the 5’ untranslated regions, where these structures participate in ribosome binding and translation (Babendure et al., 2006; Matoulkova et al., 2012). Across all 27,025 structured genes, >85% of low MFE regions overlapped UTRs, a clear bias given that 5’ UTRs collectively account for only ~24% of the gene space. Note, however, that the minMFE of genes was lower than that of their mRNAs for 58.6% of mRNAs (**Figs 1&2**) and also that 10.2% of genes changed status from structured to unstructured when considered only as transcripts. We therefore suspected that many of the structured genes contained internal regions representing TE insertions within introns. Consistent with this hypothesis, structure across the length of genes vs mRNAs differed in the bodies of these features but not in the 5’ end (**Fig 3**).

**Figure 3:**
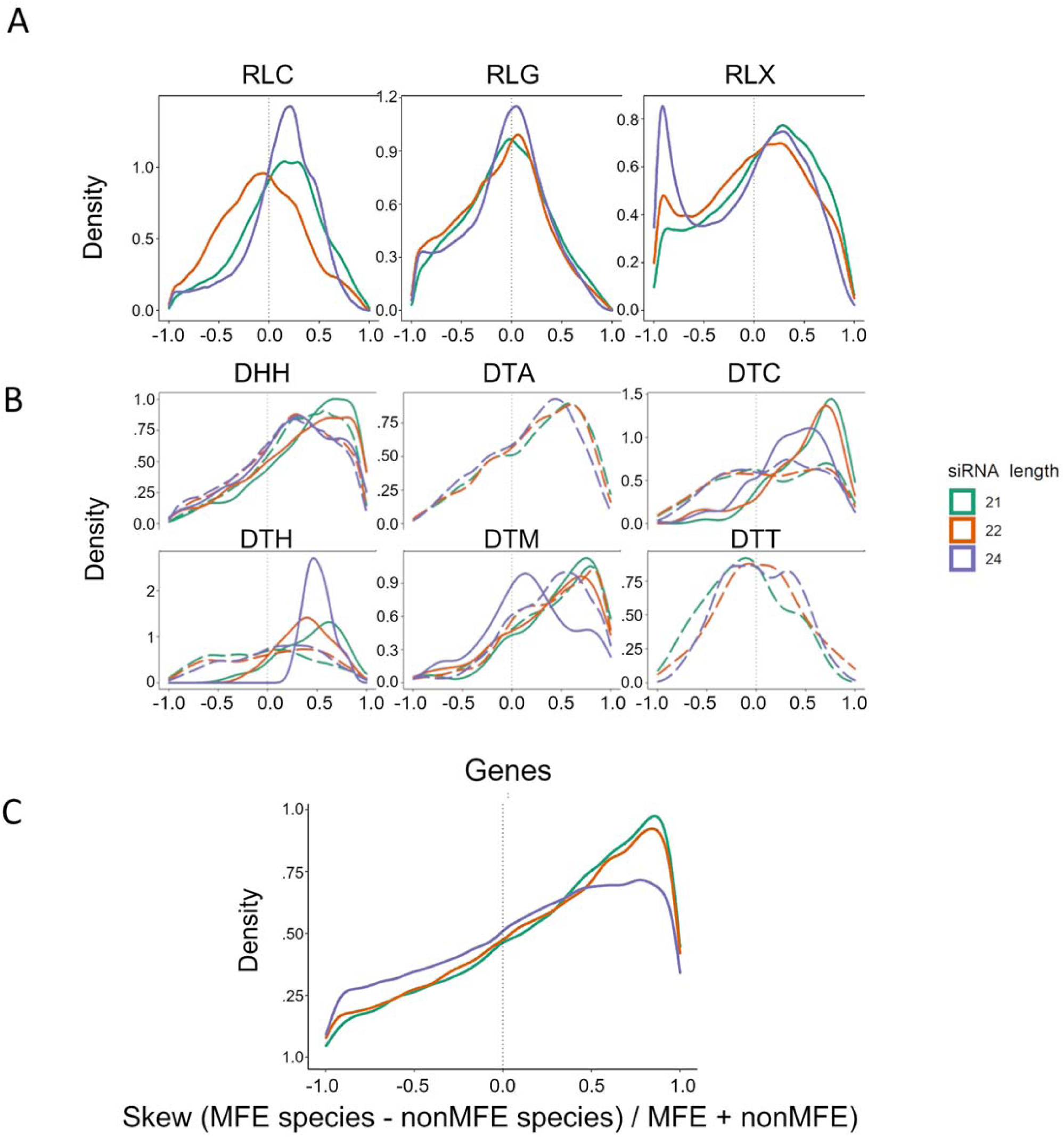
siRNA mapping skew towards structured regions. All panels use the same x-axis, which is a measure of skew [i.e., (siRNA species density in low MFE regions - siRNA species density in other regions)/total siRNA density], and the dotted vertical line represents zero where siRNA density is not skewed to either low or high MFE regions. **A.** Retrotransposons and their skew for 21, 22 and 24nt siRNAs, representing *Copia* (RLC), Ty3-Gypsy (RLG) and unknown retrotransposons (RLX). **B.** DNA transposons, with names for the three letter codes provided in Table 1. The solid lines represent autonomous elements, while dashed lines represent non-autonomous elements. **C.** Skew measured in genes.

### Strong correlations between secondary structure and siRNA counts

Under the self-priming model, we hypothesized that genomic regions of high secondary structure correlate with regions that have homology to more siRNAs. To test the hypothesis, we mapped siRNAs from 24 small RNA libraries (see **Methods**; **Table S1**) to the B73 maize genome, and then counted the number of unique siRNA sequences (i.e., distinct siRNA species) (Bousios et al., 2017) that mapped with 100% identity to genomic regions. Because of their distinct functions and origins, we separated siRNAs into three size classes (21, 22, and 24 nt).

We first examined the relationship between secondary structure and siRNAs using a linear model across all 373,485 features. The correlation coefficient was generally small - e.g., R^2^ was ~0.1 for models incorporating minMFE - but it was highly significant across all siRNA lengths for all summary metrics (**Table 2**). Extending this approach to the 15 individual feature categories and three siRNA lengths, the minMFE of a feature and the number of siRNA species per nucleotide was significant (linear model, P < 0.05 FDR corrected) for 36 of 45 comparisons (**Table S2; Fig S6**). Overall, these results indicate a consistent relationship between secondary structure and the number of siRNAs that map to features. We did note, however, some interesting outliers. First, the relationship between siRNAs and any of the MFE summary statistics was not significant for miRNAs, perhaps in part reflecting small sample sizes (n=107; **Table S2**). To this end, the LINE categories also were typically not significant, despite being heavily saturated with all siRNA size classes (**Fig S6**), reflecting both small samples and little secondary structure. We also note that the linear relationship was typically higher for 21 and 22-nt siRNA than for 24-nt siRNA (**Table 2&S2**). In genes, for example, correlations between minMFE and 21-22 nt siRNAs were significant (R2 > 0.01, P < 4.120e-106), but the correlation with 24-nt siRNAs was not (R2 = 8.35e-05, P = 0.072)(**Table S2**). Overall, however, these analyses broadly supported a genome-wide association between secondary structure and siRNA mapping.

**Table 2:** Correlation value (with FDR corrected p-value in parentheses) between MFE summary statistics and numbers of siRNAs across all 373,485 features.

| Summary Metric | 21-nt siRNA | 22-nt siRNA | 24-nt siRNA |
| --- | --- | --- | --- |
| minMFE | 0.0906 (0.00) | 0.103 (0.00) | 0.0738 (0.00) |
| meanMFE | 0.0166 (0.00) | 0.00861 (0.00) | 0.00431 (5.01e-227) |
| varMFE | 0.0179 (0.00) | 0.0170 (0.00) | 0.0235 (0.00) |
| propMFE | 0.000166 (2.95e-10) | 0.000873 (2.13e-47) | 0.000967 (2.41e-52) |

We also examined the relationship between MFE and siRNA counts *within* features. To perform this analysis, we focused only on structured features (**Table 1**) and compared siRNA mapping of all low (<-40 kcal/mol) vs. all high (>-40 kcal/mol) MFE regions of the feature. In most features, the combined length of the high MFE region was substantially longer than the low MFE regions. As a result of this length disparity, there were many cases (26% of feature:small RNA library pairs) where no siRNAs mapped to low MFE regions, particularly among features that were targeted by few siRNAs overall. Additionally, there were some cases (~10% of feature:small RNA library pairs) where no siRNAs mapped to high MFE regions. We therefore compared low and high MFE regions by removing individual features with zero siRNA counts in either the low or high MFE regions. We then measured the skew of siRNA mapping towards low MFE vs unstructured regions (see **Methods**). The skew represented the proportional difference between low and high MFE regions and ranged from −1.0 to 1.0. When the skew was negative, siRNA species were more abundant in unstructured regions, but higher skews (approaching 1.0) indicated that siRNA species were skewed towards structured regions.

As expected, *Copia/RLC* elements had positive skews, reflecting the tendency for more siRNAs to map in low MFE regions (**Fig 3a**). However, this effect was only visually detectable in 24 and 21-nt siRNAs and not 22-nt siRNAs. This result was confirmed by a linear mixed effects models, because 21 and 24-nt species/NT were significantly higher (P ≅ 0) in *Copia*/RLC low MFE regions than high MFE regions, but 22-nt siRNAs were significantly more abundant in unstructured regions (P ≅ 0; **Table S3; Fig S8**). In contrast, *Ty3-Gypsy*/RLG elements had very little visible skew in any size class (**Fig 3a**), but this TE superfamily did have slightly higher siRNA species counts in low MFE regions (P ≅ 0). DNA transposons had significantly higher siRNA counts of all size classes within low MFE regions in the mixed effect models (**Table S3; Fig S8**), and siRNA species mapping clearly skewed towards low MFE regions in most superfamilies (Wilcoxon rank sum test, all *P* < 2e-10), except in *Mariner/*DTT (*P* > 0.35)(**Fig 3b**). There was, however, some variation among DNA element superfamilies: for example, *Harbinger*/DTH elements had less (but still significant; P < 2.9e-27) skew towards low MFE regions.

One source of hidden variation that could contribute to differences in secondary structure, siRNA density, and skew is autonomy. Non-autonomous DNA transposons are not transcribed, and therefore RNA secondary structure cannot drive the creation of siRNA through self-priming. To investigate, we separated DNA transposons into nonautonomous and autonomous elements using transposase homology data (Stitzer et al., 2021)(see **Methods**), and we then repeated the skew and linear model analyses. In most cases, non-autonomous elements had less siRNA skew towards low MFE regions than autonomous (**Fig 3b**). This pattern was consistent among *Helitron*/DHH, *CACTA*/DTC, and *Harbinger*/DTH elements, but not *Mutator*/DTM elements. The differences were particularly dramatic among *Helitrons*/DHH, most of which are non-autonomous in maize (Stitzer et al., 2021), for 21–22-nt siRNAs (*P* < 7.5e-31). Note that all *Mariner*/DTT elements were non-autonomous, which is probably related to their lack of secondary structure (**Fig 1**). Overall, these results are consistent with the notion that the relationship between secondary structures and siRNAs is stronger for putatively expressed TEs.

Finally, we examined genes. Unsurprisingly, genes had homology to far fewer siRNA species than most TE types – nearly 100-times less in most cases (**Fig S6**) – but siRNA species abundance was roughly equivalent between genes and their transcripts. Although genes mapped fewer siRNAs overall, they had stronger skews than any of the TE superfamilies. For example, roughly three-fold more siRNAs (of all size classes) mapped to low vs. high MFE regions in genes, compared to the 1.5- and 1.3-fold difference in *CACTA*/DTC transposons and *Copia*/RLC retrotransposons. Consistent with these observations, linear mixed effect models were significant for higher siRNA abundance in the low MFE regions of genes for all three siRNA lengths (P ≅ 0; **Table S3; Fig S8**).

### Relationships among gene expression, structure and siRNA abundance

Genes possess regions with stable RNA secondary structure (**Fig 1 & 2**), and this secondary structure coincides with the presence of siRNAs (**Fig 3c & Table S3**). Yet, genes are usually expressed, which raises the question as to whether gene secondary structure has a quantifiable relationship with gene expression. To address this question, we used previously published RNA-seq data from 23 B73 tissues across varying developmental stages (Walley et al., 2016) to compare expression in 27,025 structured versus 5,060 unstructured genes. Structured genes had significantly higher expression than unstructured genes (t-test, P < 2e-16)(**Fig 4a**), and this was true for all tissues (**Fig S9**).

**Figure 4:**
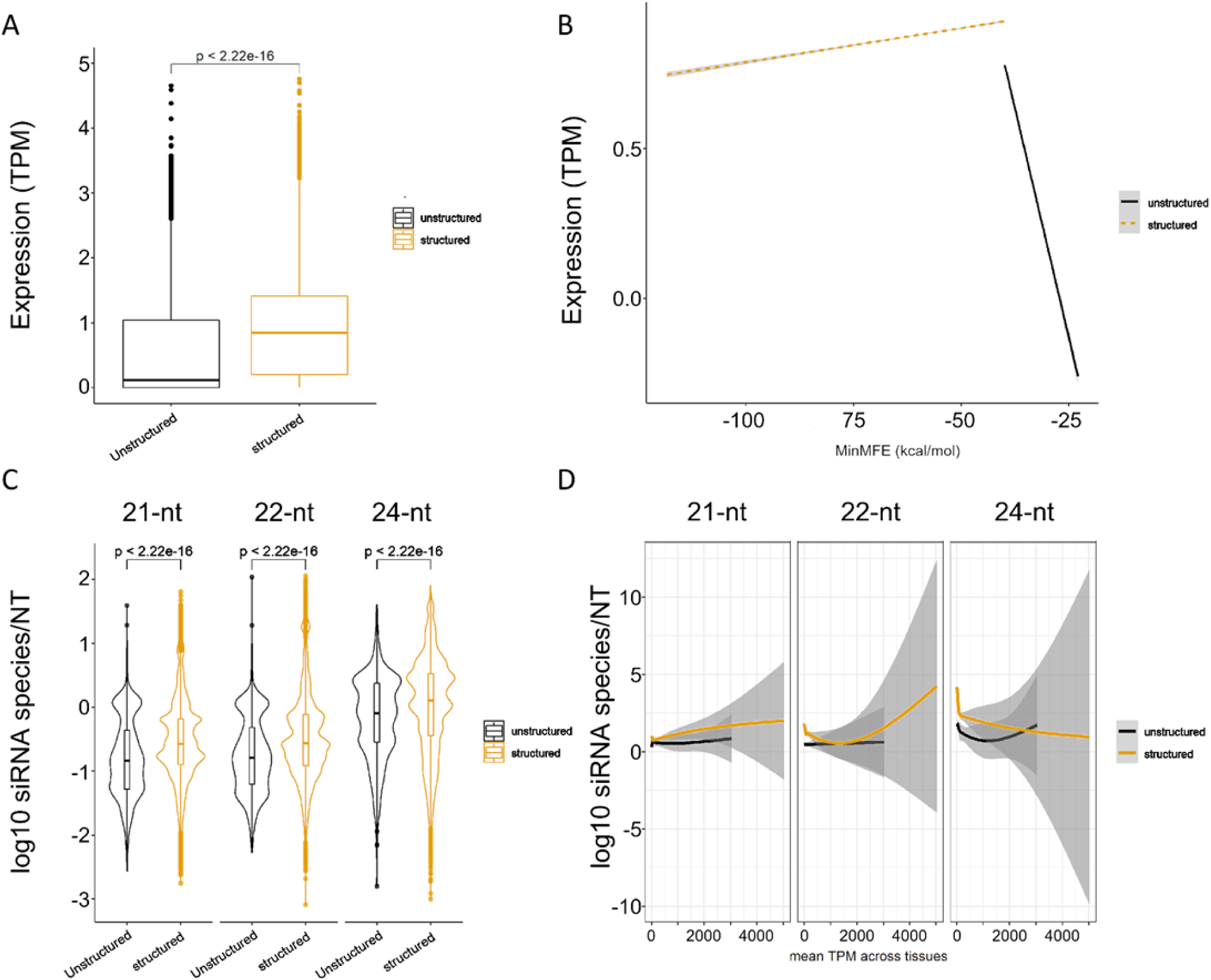
Expression between structured and unstructured genes in B73, based on data combined across 23 tissues. **A**. Difference in the overall magnitude of expression in structured vs unstructured genes. **B**. Expression as a function of minMFE for structured and unstructured genes. **C**. siRNA species per nucleotide between unstructured and structured genes. siRNA species counts from all 24 small RNA libraries were combined for this analysis. **D**. Generalized additive models of gene siRNA species per nucleotide as a function of expression, measured for both structured and unstructured genes.

We suspect, however, that many unstructured genes may be pseudogenes, since secondary structure in genes is likely essential for proper mRNA processing (Vandivier et al., 2016). We therefore also examined the effect of secondary structure as a quantitative variable among structured genes. For structured genes, minMFE correlated weakly with expression (P < 2e-16, R^2^ = 9e-4), such that genes with lower minMFE (and hence higher stability) were more lowly expressed overall (**Fig 4b**), and this was true for 16 of 23 tissues (**Fig S10**). The relationship was the opposite for unstructured genes: minMFE was highly negatively correlated with expression in every tissue (P < 2e-16, R^2^ = 0.083), so that genes with more stable secondary structure were more highly expressed (**Fig 4b**), and this was true for all tissues (**Fig S10**). We interpret these relationships as reflecting both a qualitative and a quantitative effect of secondary structure on genes. The qualitative effect is that it is important to have some structure for gene function. The quantitative effect, as reflected by the negative relationship between expression and stability, is that too much structure may not be a positive attribute for expression, perhaps due to creation of siRNAs (**Table S2**; **Fig 4c**).

We also tested whether genic secondary structure affected the relationship between gene expression and siRNA abundance. Our rationale was that higher expression leads to more opportunity for the production of siRNAs through self-priming. To investigate, we repeated the previous linear model analysis but split genes into structured and unstructured categories. The expression level of unstructured genes was not correlated with siRNA production in any size class (minimum P = 0.173). For structured genes, gene expression was uncorrelated with the number of 21-nt siRNA abundance, weakly negatively correlated with 22-nt siRNAs, and more strongly negatively correlated with 24-nt siRNAs (P = 1.284e-14, R^2^ = 1.5e-3). Given the low R^2^ values of these linear models, we represented these relationships as generalized additive models (**Fig 5d**). From these models, 21-22-nt siRNA abundance declined sharply with expression of structured genes from 0 to ~100 TPM, but the relationship became positive at higher expression levels, suggesting the possibility that higher expression leads to more siRNA production.

**Figure 5:**
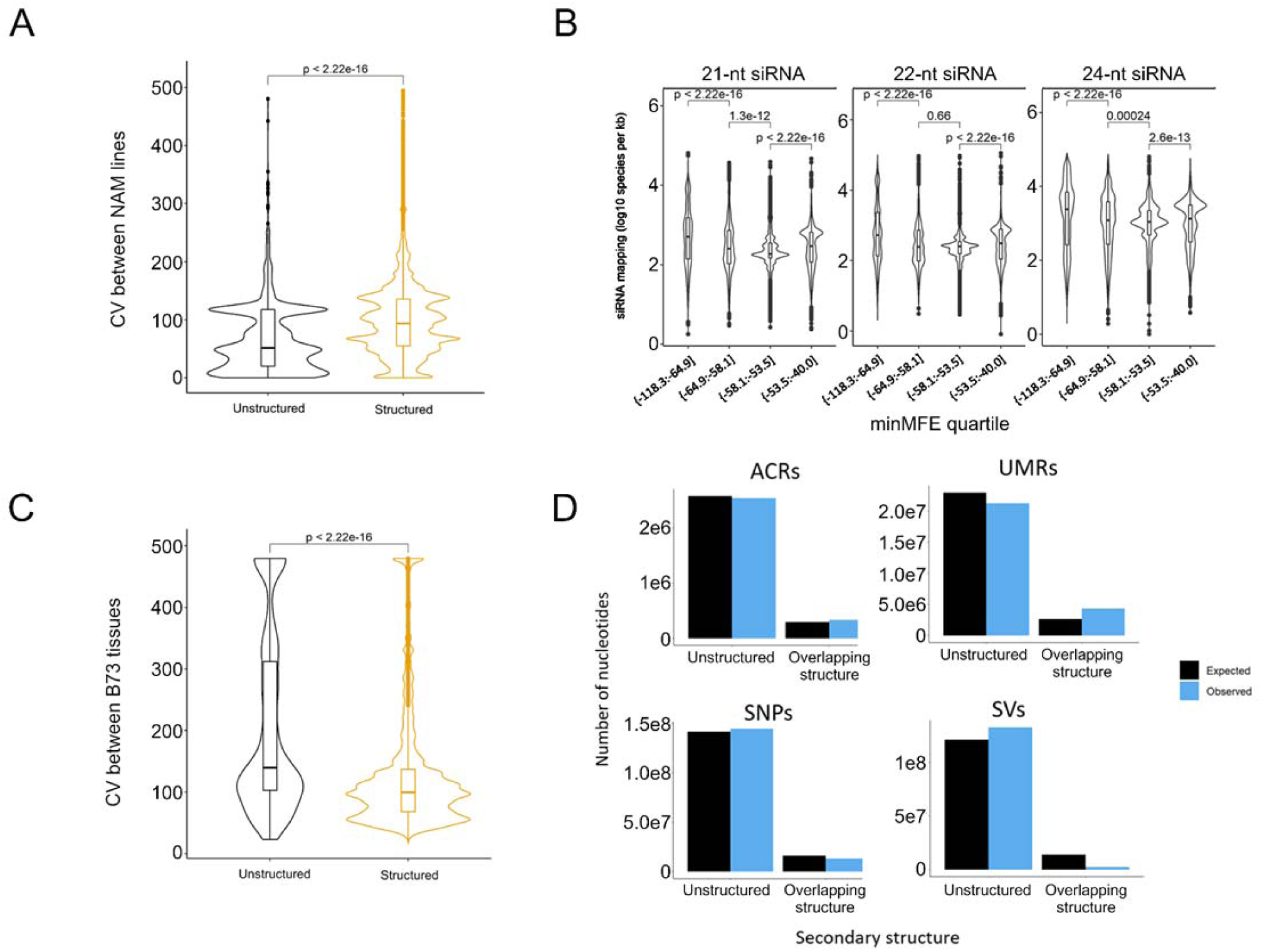
Underlying genetic and epigenetic features of structured regions across 26 inbred lines. **A**. Coefficient of variation in expression between 26 lines in structured vs unstructured genes. **B**. siRNA species mapping density between minMFE quartiles of structured genes. P-values represent outcomes of unpaired t-tests. **C**. Coefficient of variation in expression across B73 tissues in structured vs unstructured genes. **D**. Epigenetic and genetic features in low MFE and high MFE regions of genes. ACRs = Bar charts represent the overlap, in nucleotides, between epigenetic features and low MFE regions in the B73 v5 assembly. ACRs = accessible chromatin regions, UMRs = unmethylated regions, SVs = structural variants (or indels), and SNPs = single nucleotide polymorphisms. In none of the four cases did the expected and observed number of nucleotides differ significantly between structured and unstructured regions.

### Structured genes have higher expression variance among maize inbred lines

The complex relationships among gene expression, siRNA abundance, and secondary structure could represent a trade-off between the essential functions of genic secondary structure and the potential deleterious effects of self-priming and hairpin siRNAs. If this trade-off occurs, one expects that siRNAs modulate expression of genes. We tested this idea by gauging the relationship between secondary structure and gene expression across the 26 nested association mapping (NAM) founder lines (McMullen et al., 2009), which have corresponding genomic, expression and epigenetic data (Hufford et al., 2021). We predicted that genes with high secondary structure and matching siRNAs are more variable in expression due to the hypothesized trade-off.

For these analyses, we assumed that the secondary structure designations from B73 (structured or unstructured) applied to the 28,866 shared genes across the 26 lines (Hufford et al., 2021). With these designations, we first confirmed that structured genes were more highly expressed across all lines using pangene expression data (Hufford et al., 2021)(**Fig S11**; unpaired t-tests, all P < 2.2e-16), indicating that this subset of genes broadly shared the patterns of expression of B73 across lines. We then measured gene expression among lines. For each shared gene, we calculated the coefficient of variation (CV) of expression across the 26 lines. Structured genes had a significantly higher CV than non-structured genes (P < 0.01, permutation test)(**Fig 5a**). This was true both for comparisons between all genes in each group and between a downsampled subset of structured genes that was equal in size to the set of unstructured genes.

One obvious difference between these subsets of genes is the magnitude of their expression; although CV is standardized by the mean, more highly expressed genes may also be more variably expressed. To examine this possibility more thoroughly, we fitted a linear model of expression CV as a function of B73 gene expression, but there was a weak negative correlation (R^2^ = 6.349e-05, P = 0.046, estimate = −0.01). We thus believe that higher variability in structured genes is not simply a product of them being more highly expressed.

### Potential causes of variable expression of structured genes

The higher CV for expression of structured genes raises a further question: Is this variation consistent with the self-priming model - i.e., these genes are more variable across lines because of the stochasticity associated with self-priming and some modulation of expression via siRNA production? Under this hypothesis, more siRNAs lead to more expression variation across lines, due to the stochastic effects of RNAi-like dynamics. To investigate this possibility, we fit a linear model of expression CV as a function of siRNA density and found that CV was positively correlated with siRNA abundance (P = 6.7e-283; R^2^ = 0.010). To see if an effect was discernible between structured genes of variable minMFE values (as suggested by **Fig 4b**), we separated structured genes into four quartiles based on their minMFE and then plotted the number of siRNAs that map to each gene in B73. Consistent with our hypothesis, genes in the lowest minMFE quartile mapped more siRNAs than the other three quartiles for all three siRNA lengths (**Figure 5b**), and minMFE was significantly correlated with CV in a linear model (P = 5.8e-79; R^2^ = 3.1e-3).

The above evidence suggests that higher CVs for expression are related to the number of siRNAs that map to a gene, but how? One possibility is that self-priming of structured genes drives RdDM and chromatin closure. To test this idea, we used ATAC-seq and methyl-seq data (Hufford et al., 2021) to assess variability of accessible chromatin regions (ACRs) and unmethylated regions (UMRs) among lines (see Methods). For each line, we identified whether variable epigenetic states (in ACRs and UMRs) overlapped with low MFE (< −40) regions, expecting that ACRs/UMRs overlapping low MFE regions were less abundant than expected and more rarely conserved between lines. We found that ACRs and UMRs overlapped low MFE regions more frequently than expected by random chance, but not significantly so (**Fig 5d**). Taken together, these results suggest regions of secondary structure rarely vary due to epigenetic marks and these marks are not the cause of high CV among structured genes.

We next assessed whether expression is disrupted by transcript degradation, as we might expect under RNAi-like dynamics. We reasoned that stochastic fluctuations due to transcript degradation should be visible both within and between lines, if it is the process that drives expression CV. Returning to the B73 expression data across 23 tissues (Walley et al., 2016), we measured the CV of structured vs unstructured genes, this time between tissues rather than between lineages. In this case, CV was significantly lower in structured genes than in unstructured genes (**Fig 5c**), and so we could not detect the expected effect across B73 tissues.

A final possibility is that genetic mutations, including structural variants (SVs) and SNPs, occur more often in structured regions. Our motivations for this idea were that highly structured regions may have different mutation rates (Hoede et al., 2006; Weigel et al., 2022) and also that indels occur commonly in the palindromic regions of Sireviruses (Bousios et al. 2016). To perform these analyses, we used existing SNP and SV calls from the NAM lines (Hufford et al., 2021), again assuming a structured gene in B73 was also structured for the remaining lines. For deletions, we compared their % overlap with B73 structured regions; for SNPs, we measured the proportion that fell within B73 structured regions. We found that SNPs occured at roughly their expected frequencies in structured regions, but indels were highly depleted in the structured regions of genes (**Fig 5d**). Although these dynamics are likely mediated by purifying selection, structured regions do not appear prone to particularly high rates of mutation. Hence, high mutation rates also do not adequately explain the elevated CV for the expression of structured genes.

## DISCUSSION

We have profiled secondary structure in annotated maize features, including TEs and genes, to establish a general relationship between RNA-folding and siRNA-mapping. To our knowledge, our study is the first to compare secondary structure characteristics among TE superfamilies, revealing some striking qualitative differences. We find, for example, that LTR retrotransposons and helitrons have regions of strong secondary structures, that SINEs and LINEs have very little secondary structure, that TIR elements vary in structure according to superfamily (**Fig 1**), and that secondary structure and siRNA dynamics differ between autonomous and non-autonomous TEs (**Fig 3**). Similar to previous work in *Arabidopsis* (Li et al., 2012), we have also detected structured regions within maize genes, particularly within 5’ UTRs. These results are not unexpected, because 5’ stem-loop structures are critical for translation (Vandivier et al., 2016). However, we have also shown that genes with detectable structures have higher expression than those without structure (**Fig 4a**), more abundant mapping by siRNAs to structured regions (**Fig 3**), and higher coefficients of variation in gene expression across individuals (**Fig 5a**). Taken together, our observations provide a foundation for interpreting the impact of secondary structure on genome function and evolution.

### Secondary structure and siRNA production

For detecting secondary structure, we have included two positive controls: miRNA precursor loci (Wang et al, 2009) and *Copia*/RLC elements (Bousios et al., 2016). As expected, these two feature sets have the most extreme MFE statistics among our sample. *Copia*/RLC elements have the lowest minMFE values, reflecting previously recognized regions of strong secondary structure (**Fig 1a**), but it is worth noting that other retrotransposons have similarly low minMFE distributions (**Fig. 1a**). miRNAs do not have the lowest minMFE among our dataset, perhaps due to their short length, but they have the lowest meanMFE and the highest proportion of their sequences included in low (<-40 kcal/mol) MFE windows (**Fig 1**). These two positive controls provide a basis for comparison to other feature types. For example, across the remaining categories of TEs and genes, DNA elements generally have less structure than retrotransposons (**Fig. 1**); in fact, we have detected no LINE elements with significant structure relative to randomized sequences (**Table 1**). Another consistent thread is that most (69%) genes have evidence for higher-than-expected local secondary structure in their 5’ UTRs, with lower minMFEs and mean MFEs than miRNAs and some TE types (**Fig. 1**).

One must be careful with these comparisons in light of the limitations of our methods. We have relied on RNAfold predictions applied to small overlapping windows. We chose this approach because it has precedence in the literature (Wang et al., 2009; Bousios et al., 2016) and also because having identical window sizes provides a basis for comparing properties within and between features. However, we recognize that bioinformatic predictions of secondary structure often do not correspond to *in vivo* assessments (Wang et al., 2013) and also that we have simplified secondary structure to quantitative summaries. We thus recognize that our secondary structure inferences are approximations.

The more important question is whether our approximations bias our results, particularly our finding that secondary structure commonly correlates with siRNA mapping. We do know there is bias in some of our summaries - e.g., minMFE is correlated with feature length and low MFE regions are more likely in sequences with high G:C composition. Although we have tried to control for these biases (e.g., by using multiple summary statistics and randomizing primary sequence), some undoubtedly remain. Moreover, if secondary structures are functional, we suspect that combining non-functional with functional TEs probably under-represents secondary structure among active TEs, perhaps obscuring signals of consistent secondary structure in specific TE regions. Our data also have a high false negative rate, given that we detect that most (71%), but not all, pre-miRNA loci are structured based on minMFE values and sequence randomization (**Table 1**). Overall, these considerations suggest that our analyses underestimate, rather than overestimate, the prevalence of secondary structure. However, even with this likely underestimation, we detect consistent negative correlations between MFE metrics and siRNA diversity (**Table 2&S2**). These correlations are significant across features and also between low and high MFE regions within features (**Fig 3**). It is highly unlikely that our mapping of secondary structure drives these correlations, because error in secondary structure measurements should lead to weaker or non-existent correlations. Since both genes and TEs exhibit this relationship, we conclude that the structure:siRNA correlation is a general characteristic of the maize epigenome.

Given known pathways of miRNA biogenesis (O’Brien et al., 2018), we believe the most likely explanation for the observed pattern is self-priming - i.e., that dsRNA loops lead to siRNA production. This conclusion is bolstered by the fact that we detect this relationship in each feature category (**Table S2**), and also by our observation that siRNA skew is most pronounced for putatively expressed genomic regions – like genes and autonomous (vs. non-autonomous) elements (**Fig. 3**) – where we expect self-priming to be most active. It is important, however, to consider other explanations. For example, it is possible, although we believe unlikely, that the siRNA:structure correlation is due to a biological mechanism other than self-priming. In Arabidopsis, miRNA target sites within mRNAs are significantly less structured than surrounding regions (Li et al., 2012), which is thought to confer accessibility to the endoribonucleases involved in RNAi (Vandivier et al., 2016). This pattern hints that small RNA binding and RNAi is less effective in structured regions of TEs than in non-structured regions, as is likely the case in viruses (Gebert et al., 2019). If this is the case, it is possible that the structured regions of TEs include sequences that are important for TE function due to their primary sequence rather than their secondary structure and, further, that the secondary structure has evolved to protect those primary sequences from targeting through RNAi. In this explanation, the structured regions are first highly targeted by siRNAs and then structure evolves as a component of the evolutionary arms race between TEs and their hosts.

Whatever the cause, we find the structure:siRNA relationship to be consistently significant across genomic features. If self-priming is the primary mechanism, our results offer additional insights into the process. First, we note that correlations between siRNAs and secondary structure explain a relatively low proportion of the variation in siRNA mapping across the genome. For example, across the entire dataset of 373,485 features, minMFE explains at most 10% of the siRNA mapping results (**Table 2**). This value can be higher within other MFE metrics and specific feature categories - e.g., propMFE explains 24% of siRNA variation in *CACTA*/DTC elements - but the explanatory power of MFE statistics is typically between 3.8% (the median R2 across feature types for varMFE) to 10% (the median for meanMFE) (**Table S2**). These R2 values are consistent with the fact that self-priming is only one of several mechanisms that produce siRNAs (Carthew & Sontheimer, 2010), and they provide an estimate of the contribution of self-priming relative to other mechanisms of siRNA production. Second, we also infer from these data that miRNA-like secondary structures are probably insufficient to lead to TE silencing on their own. For example, we have observed that *Mutator*/DTMs provide substantial evidence for strong secondary structures (**Fig. 1**), but we also know that a separate silencing element (*Mu-killer*) is required to initiate silencing of an active element (Slotkin et al., 2003). We also know that epigenetic marks like methylation often spread along sequences. TE silencing is tightly maintained because, after the initiation of RdDM, methylation (and siRNA generation) spreads from the initially targeted locus to the length of the entire TE (Ahmed et al., 2011). This spreading suggests that self-priming alone is not sufficient for a complete silencing response, even if self-priming initiates the response. This pattern differs remarkably from expressed genes, where either silencing does not spread across the gene length or it is removed after it is deposited (Gong et al., 2002; Penterman et al., 2007). We emphasize that although positive skew (towards +1) indicates higher siRNA abundance in structured regions and may result from self-priming, a lack of skew does not imply that the *initial* silencing of an element did not result from self-priming.

### The self-priming trade-off

The potential for self-priming in genes has been previously suggested by Li et al. (2012), who found that mRNA transcripts with more stable secondary structure corresponded to higher small RNA expression and lower expression in *Arabidopsis*. We have built on this work in three ways: first, we have extended the siRNA:structure relationship to maize, which has a much larger, more TE-rich genome. Given that the strength of TE-silencing machinery may differ between large and small genomes (Hollister et al., 2011), it is notable that gene self-priming/RNAi is shared between maize and *Arabidopsis*. Second, we have shown that secondary structure does not universally negatively correlate with gene expression. Rather, the relationship is tiered. There is a qualitative difference in expression between genes with and without secondary structure (**Fig 4A&B**), reflecting that it is critically necessary for some aspects of gene function. However, the stability of secondary structure in significantly structured genes correlates negatively with expression (**Fig. 4B**), suggesting that there can be, in fact, “too much of a good thing” when it comes to secondary structure. The potential functional consequence of “too much” is illustrated across NAM lines, because structured genes with higher coefficients of variation tend to map more siRNAs (**Fig. 5B**).

These results provide evidence of an evolutionary tradeoff between selection for stable secondary structure against too much secondary structure. In genes, this tradeoff heavily favors the presence of secondary structure, because most genes (69%) have significant structure with 87% having miRNA-like minMFE values. Given the strong functional need for secondary structure in genic transcripts, the trade-off makes sense for genes, but two questions remain. First, if self-priming does occur, as our data seem to suggest, then why are genes not silenced by the diverse array of mechanisms that act on TEs? We began this study predicting that genes and TEs differed in their secondary structures if they act as potential signals for silencing. Not only have we not detected an obvious difference, but we have found some unexpected evidence for self-priming in genes as measured by siRNA skew (**Fig 3C**). So, why are genes not silenced? We do not have a complete answer, but we believe it must rely on the bevy of differences between hetero- and euchromatin. It is known, for example, that genic regions have distinct sets of chromatin markers relative to heterochromatin and also that demethylases like *Increased in Bonsai Methylation 1 (IBM1)* and *repressor of silencing 1* (ROS1) (Gong et al., 2002; Penterman et al., 2007) actively demethylate expressed genes (Saze et al. 2008; Miura et al. 2009). Some aspects of genic methylation are under selection (Muyle et al., 2022), and selection will be particularly strong against mechanisms that silence genic regions. We hypothesize that these mechanisms have evolved in part to counter the potentially deleterious effects of self-priming.

The second question focuses on TEs: if secondary structure leads to self-priming and the potential initiation of the silencing cascade, why have TEs not evolved to lack secondary structure? By doing so, they could in theory escape one component of the host response. We cannot answer this question definitively either, but we suspect it again revolves around the potential function of secondary structure. In Sireviruses (represented principally by the *copia*/RLC elements in this study), evidence suggests that the palindromic regions act as a *cis*-regulatory cassette (Grandbastien et al., 2015). In fact, studies of different TE families in different organisms have revealed that *cis*-regulatory regions are often arranged as arrays of complex, sometimes palindromic, repeats (Vernhettes et al., 1998; Araujo et al., 2001; Fablet et al., 2007; Ianc et al., 2014; Martinez et al., 2016), again suggesting that secondary structure often assumes a *cis*-regulatory function. Another consideration is that retrotransposons and autonomous DNA elements need to replicate by expressing and translating their genes by co-opting the host’s translation machinery. This suggests that secondary structure is likely to be as crucial for some aspects of the TE life-cycle function as it is for genes. This idea explains the prevalence of secondary structure in retrotransposons (**Fig. 1**) and the exaggerated siRNA skew in autonomous vs. non-autonomous TEs (**Fig. 3**).

Overall, our results show that stable secondary structures - as inferred from low MFEs - correlates positively with siRNA abundance and may be shaped by a trade-off between functional requirements and the potential disruptive effects of self-priming. We hope this work sparks further exploration of the roles of secondary structure in plant genome evolution, including the population genetics of mutations in regions of secondary structure (Ferrero-Serrano et al., 2022), their roles in the stress response (Zhu et al., 2018), and comparative analyses of secondary structure characteristics between species.

## METHODS

### B73 annotation and secondary structure prediction

Version 4 of the B73 maize genome and version 4.39 of the genome annotation were downloaded from Gramene (www.gramene.org). B73 TE annotations (B73v4.TE.filtered.gff3.gz) were retrieved from https://mcstitzer.github.io/maize_TEs/ (Jiao et al., 2017). TE and gene annotations were cleaned for redundancy (e.g., the same feature annotated by different annotation authorities) using custom scripts, and separated into annotation files for different feature categories. Bed files were then generated for each annotation feature, with a standardized naming convention for each feature: Feature Type::Chromosome:Start Position-End Position (e.g., exon::Chr1:47261-47045).

Fasta files for each feature were generated using Bedtools getFasta. These fasta files were divided into 110 nucleotide sliding windows (1-nt step size) for use in the secondary structure prediction program RNAfold v2.4.9 from ViennaRNA (Hofacker et al., 2011). MFE calculations per window were extracted from RNAfold predictions using a Python script, and the MFE summary metrics (minMFE, meanMFE, varMFE, and propMFE) were calculated for each feature, based on all windows in that feature. As described in the main text, minMFE was calculated as the lowest MFE window in the feature; propMFE was calculated as the number of low MFE windows (<-40 kcal/mol) divided by the total number of windows in the feature; meanMFE was the mean of all 110 bp window MFE values; and varMFE was the variance across all windows. Bed files representing regions of significant, miRNA-like secondary structure were created by combining all overlapping windows of <-40 kcal/mol MFE. Overlapping MFE windows were converted to bed format using an inhouse Python script. The location, category and summary statistics for each feature is available as **Table S5**.

To determine whether a feature contained significant structure, the feature sequence was randomized by shuffling the position of nucleotides across the length of the feature. This approach maintained the GC content of the feature but not the primary sequence. Randomized sequences were then subjected to identical MFE calculations - i.e., they were split into 110 bp windows for RNAfold prediction. This process was repeated five times for each feature, and the minimum MFE of each randomization was recorded. The significance of observed structure vs the five randomizations was assigned using a Wilcoxon one-sided test with Benjamini-Hochberg correction in R.

For plotting the location of low MFE regions across featureS (**Figs 2 & S4**), we split each feature into 100 equally-sized bins across the length of the feature from 5’ to 3’ end and counted the number of <-40 kcal/mol regions overlapping each bin. To find motifs in low MFE regions of different feature types, bed files from concatenated low MFE regions were extracted using Bedtools getfasta. These fasta files were fed into the MEME motif finder (v5.4.0)(Bailey & Elkan 1994) with the DNA alphabet in Classic mode (i.e., enrichment of sequences in a single reference sequence and no control sequence) for each feature category. We selected the top 10 overrepresented sequences.

The scripts used for MFE calculations and analyses are available on Github (https://github.com/GautLab/maize_te_structure).

### Small RNA Library Analysis

Small RNAseq libraries were downloaded using NCBI SRA tools and SRAExplorer (https://github.com/ewels/sra-explorer), from the sources indicated in **Table S1**. Adapters, regions with low quality, and low quality reads were trimmed from small RNA RNAseq libraries using fastqc and cutadapt v0.39 (Bolger et al., 2014). Adapter sequences varied among libraries, and so were identified and validated in each library using a custom bash script that searched for sets of known maize siRNAs of each length (21–24 nt) in each unprocessed library and confirmed the identity of the adapter sequence connected to each known siRNA sequence. The list of adapters derived for each library is included in **Table S4**. Trimmed reads were then filtered and split based on size matching 21, 22 and 24 nucleotides in length, creating three fastq files for each library. We considered all small RNAs of these sizes to be “siRNA” irrespective of their sequences. We identified the unique siRNA sequences, which we refer to as ‘species’, following previous methods (Bousios et al., 2016, 2017).

siRNA species were mapped using Bowtie2 v2.4.2 (Langmead & Salzberg 2012) to the B73 genome, preserving only perfect alignments. Samtools v1.10 (Danecek et al., 2021) was used to convert and sort the alignment output. Bedtools bamtobed was used to convert the sorted BAM file to bed files. siRNAs from each library were mapped separately for all three lengths, generating a total of 72 (3 sizes × 24 libraries) alignment files. Both uniquely and non-uniquely mapping siRNAs were used to calculate the number of siRNA species corresponding to each genomic locus (https://pubmed.ncbi.nlm.nih.gov/28228849/), and strand was not taken into account. Thus, any given position in the genome can be overlapped by several siRNA species, up to two-times the length of the siRNA size class in question (21, 22, or 24).

Bedtools was used to find intersections and coverage counts (per nucleotide) between the siRNA alignment bed files for each library and the MFE region bed files. Subsequently, the siRNA alignment bed files were split into two categories: alignments that intersected low (<-40 kcal/mol) MFE regions and those that did not. Coverage and count files were subsequently generated that contained information of how many siRNA species aligned at each nucleotide, and coverage files contained a normalized count per nucleotide for classification. Normalization was performed by summing the counts and dividing by the length of the region in nucleotides across all internal positions from 0 to n; 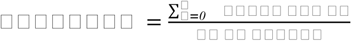.

For correlations between siRNA species density vs. MFE measurements of features (**Table 2**), linear models of siRNA species per nucleotide as a function of secondary structure metrics (minMFE, meanMFE, etc) were fitted using the base R (v4.1.0) lm() function. To fit these models, siRNA species were summed across all 24 libraries for each feature so that observed siRNA species had an equal weight across libraries. These linear models can be expressed as:

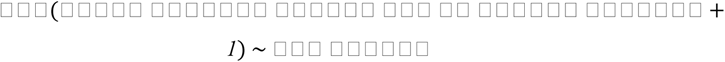

To test the significance of differences in siRNA species density between high and low MFE regions within features, mixed effects models were fit for each siRNA size class using the R package *lme4* (Bates et al., 2015). In these models, siRNA mapping counts from each library were **not** combined, meaning that each smRNA library:feature pair was counted individually. These mixed effects models can be expressed as:

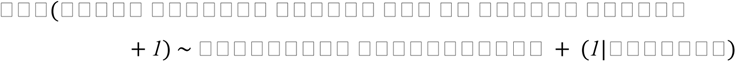

Skew measurements (**Fig 4**) were calculated separately for each TE superfamily and genes as 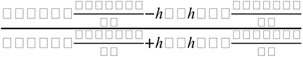. For these calculations, feature-library pairs with zero siRNA species in either non-structured or structured regions were removed from each dataset. We further tested skew differences from zero using Wilcoxon one-sided tests in R.

Autonomous vs non-autonomous designations for TEs were defined differently depending on TE type, but they were determined based on the presence or absence of open reading frames within the TEs, as identified by Stitzer et al. 2021 (downloaded from https://github.com/mcstitzer/maize_genomic_ecosystem). TIRs were considered autonomous if they contained sequence homology to a transposase, and helitrons were considered autonomous if they contained *Rep*/*Hel,* as per Stitzer et al. (2021).

### B73 RNAseq analyses

B73 gene expression data was downloaded from the ATLAS expression database (www.ebi.ac.uk/gxa/) in transcripts per million (TPM) based on RNAseq data from 23 maize tissues (E-GEOD-50191)(Walley et al., 2016). The statistical significance of differences between expression of genes in different structure classifications was determined using unpaired t-tests between structured and unstructured genes. Linear models of expression versus each measurement of secondary structure were separately fit for expression in each tissue type and graphed using ggplot2 (). These linear models can be expressed as:

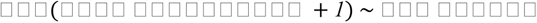

### Comparative analyses between NAM lines

Expression, ATAC-seq, methylation, and SV data for each NAM line were downloaded with B73 coordinates from CyVerse at https://datacommons.cyverse.org/browse/iplant/home/shared/NAM/NAM_genome_and_annotati on_Jan2021_release (Hufford et al., 2021). Secondary structure predictions were performed in B73 assembly V4, so gene IDs were converted to V5 using the EnsemblPlants ID History Converter web tool (https://plants.ensembl.org/Zea_mays/Tools/IDMapper). Coordinates of TEs and structured regions were converted using the EnsemblPlants CrossMap converter (https://plants.ensembl.org/Zea_mays/Tools/AssemblyConverter) with the B73_RefGen_v4 to Zm-B73-REFERENCE-NAM-5.0 parameter.

Normalized expression data were downloaded in RPKM format from merged RNAseq libraries from CyVerse at https://datacommons.cyverse.org/browse/iplant/home/shared/NAM/NAM_genome_and_annotati on_Jan2021_release/SUPPLEMENTAL_DATA/pangene-files. Only data from genes shared among all lines (as determined by Hufford et al.) were included. These data include RNAseq normalized across eight tissues in each line: primary root and coleoptile at six days after planting, base of the 10th leaf, middle of the 10th leaf, tip of the 10th leaf at the Vegetative 11 growth stage, meiotic tassel and immature ear at the V18 growth stage, anthers at the Reproductive 1 growth stage. Details for how these data were normalized can be found in Hufford et al., 2021.

The coefficient of variation (CV) of expression was calculated for each gene between the 26 lines using the normalized RPKM expression data from Hufford et al., 2021, using functions in base R. We plotted CVs between categories of structure (structured and unstructured) using ggplot2 () and determined statistical significance of differences between categories using unpaired t-tests in R. We measured these differences in two different ways: first, using all genes and, second, removing genes with CV = 0 (920 genes, 3.3% of genes). We also built a linear model with *lm*() in R to correlate the magnitude of gene expression in B73 with the CV of that gene across lines. This linear model can be expressed as:

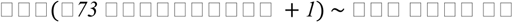

We also measured epigenetic and genetic features across the NAM lines. For the former, we concatenated ACRs and UMRs that overlap positions between lines, producing a set of merged ACRs and UMRs. We produced these merged sets using the R libraries IRanges and GenomicRanges (Lawrence et al., 2013). We kept track of the number of NAM lines that overlapped to produce a merged ACR and its position in B73. In contrast to epigenetic features, which we assumed had the potential for similar functions even when they overlapped even if the exact positions did not match, structural variants (SVs) were only counted as shared between lines when exact coordinates matched. The expected overlap was calculated as the proportional of genic space taken up by low MFE regions * the total length of features. Custom scripts for these analyses can be found at https://github.com/GautLab/maize_te_structure, and additional supplementary files can be found at https://figshare.com/projects/siRNAs_and_secondary_structure_in_maize_genes_and_TEs/150714.

## ACKNOWLEDGEMENTS

This work was supported by NSF grant to B.S.G. and by a Royal Society Fellowship grant to A.B.

## AUTHOR CONTRIBUTIONS

Designed the research: GTM, ES, AM, AB and BSG

Performed research: GTM, ES

Contributed new computational tools: ES

Analyzed data: GTM, ES, AM

Wrote the paper: GTM, AM, AB and BSG

---

**Figure S1:**
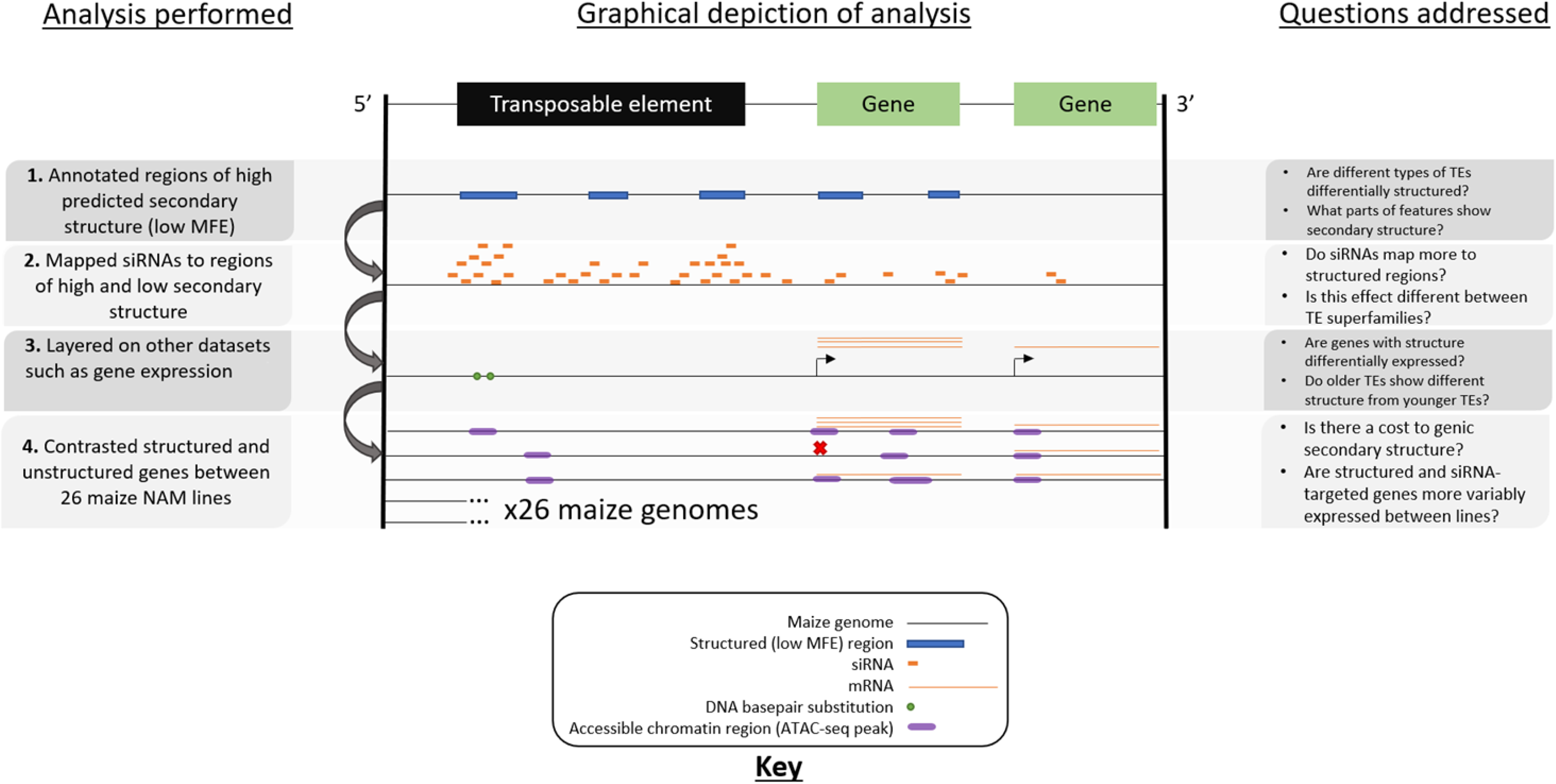
Scheme of analyses carried out. Each sequential layer includes an additional analysis performed on annotated features, including genes, ncRNA loci, and TEs. We (1.) annotated regions of predicted miRNA-like secondary structure (figs 2-3; Table 1), then (2.) mapped siRNAs across features of varying secondary structure and between regions with miRNA-like structure and without (Fig 4; Table 2), then (3.) compared expression (Fig 5), and (4.) examined underlying genetic and epigenetic features of genes with differing secondary structure between 26 inbred NAM lines representing the breadth of global maize diversity (Fig 6)

**Figure S2.**
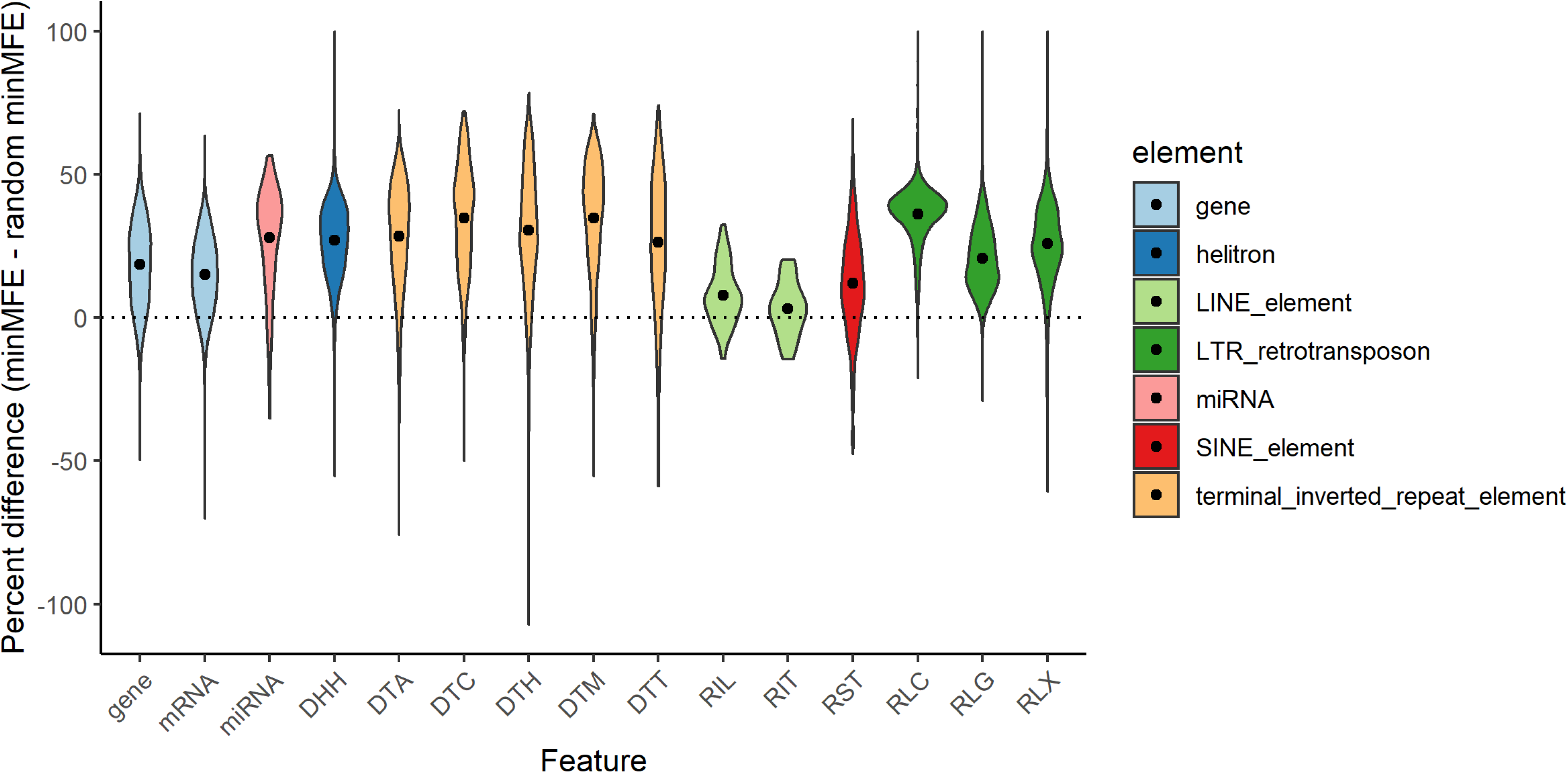
Distributions of percent differences between observed and random minMFEs in each feature type. Differences represent how much *more negative* (and therefore more stabily structured) observed minMFEs were compared to mean minMFE across five randomizations. To find percent differences, these differences were divided by the observed minMFE and multiplied by 100 [e.g., if the observed minMFE was −100 and the mean randomized minMFE was −50, percent difference would be ((−100 + −50) / −100) * 100 = 100%]. Superfamilies are colored by their broader TE category (LTR, TIR, etc.) and dots represent the mean of each distribution. The dotted line represents 0%, or zero difference from random minMFE.

**Figure S3:**
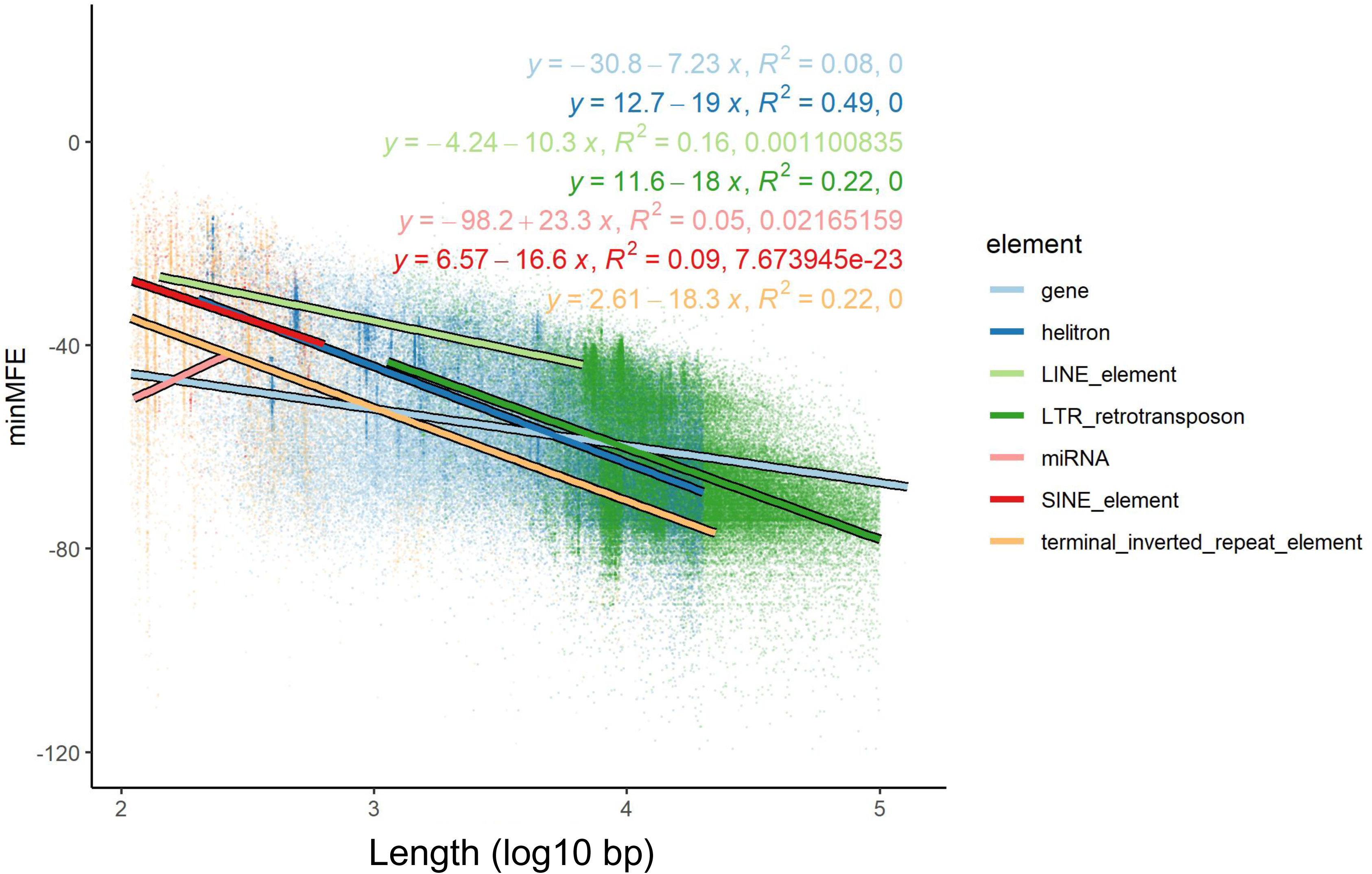
Linear models of minMFE as a function of length. Each type of TE was modeled separately using its length from 5’ to 3’ end and observed minMFE value. Plots represent simple linear models from the lm() function in R, and colored text represents the formulae, R2 value, and p-value of each regression.

**Figure S4.**
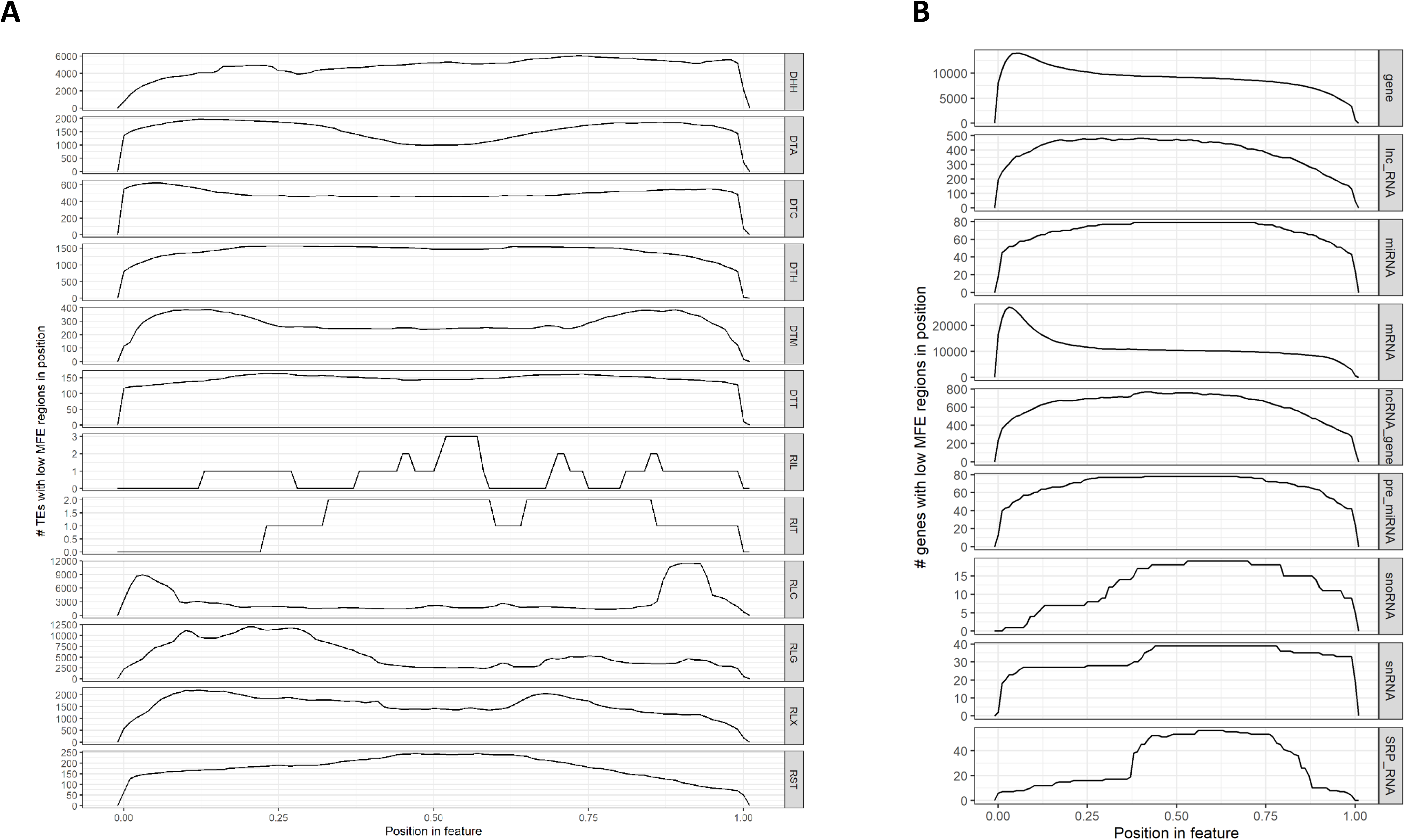
Landscapes of structured regions across feature types. (**A**) Metaprofiles across TE superfamilies, and (**B**) metaprofiles across genes and ncRNAs. Each row represents a metaprofile combining data from all members of each feature type. Features were divided into 100 equally sized bins from the 5’ end to the 3’ end, and the number of features with low MFE (<-40 kcal/mol) windows overlapping each of these bins was counted. A peak in the landscape therefore represents a region of the feature type which often shows very stable secondary structure. All rows share the same X axis, which is represented proportionally across the length of the feature from 0.00 (5’ end) to 1.00 (3’ end).

**Figure S5.**
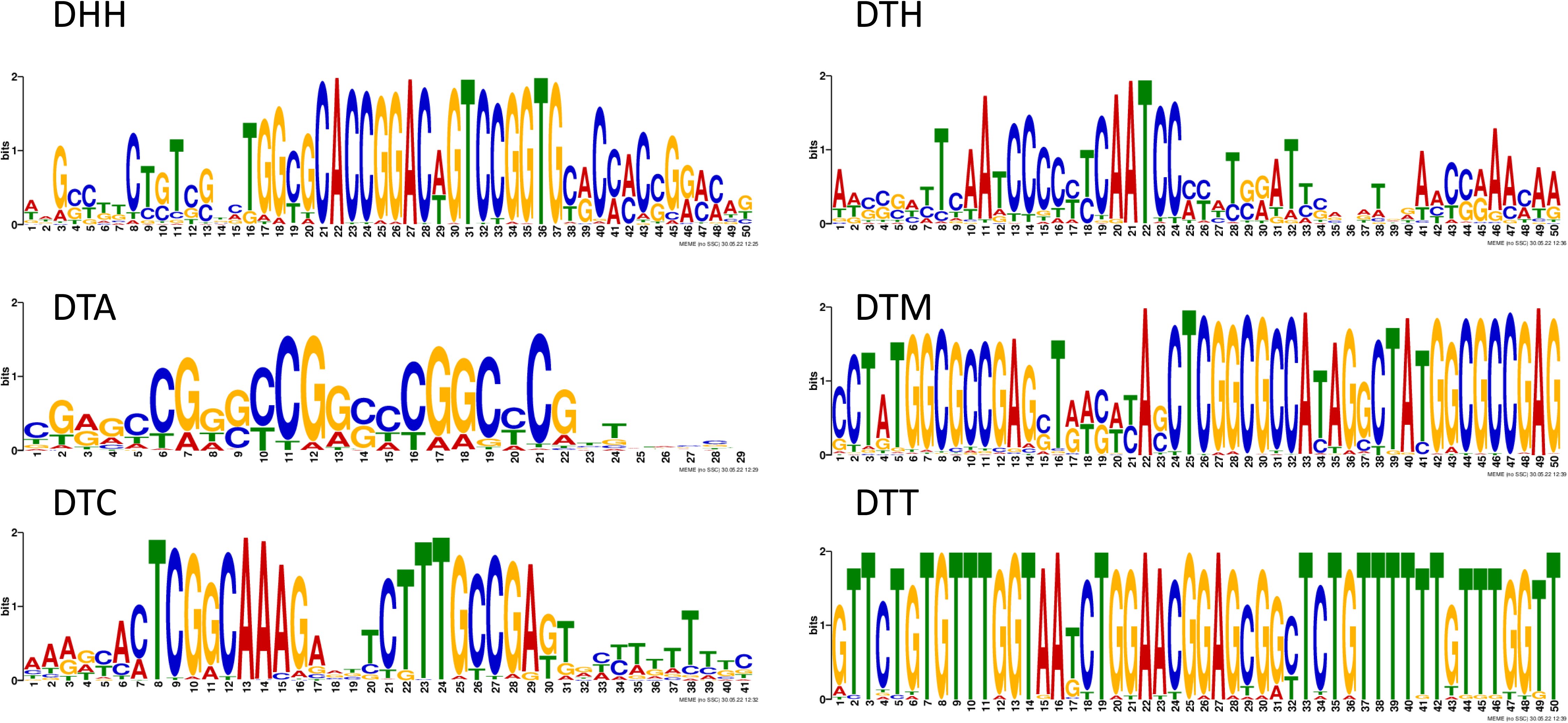
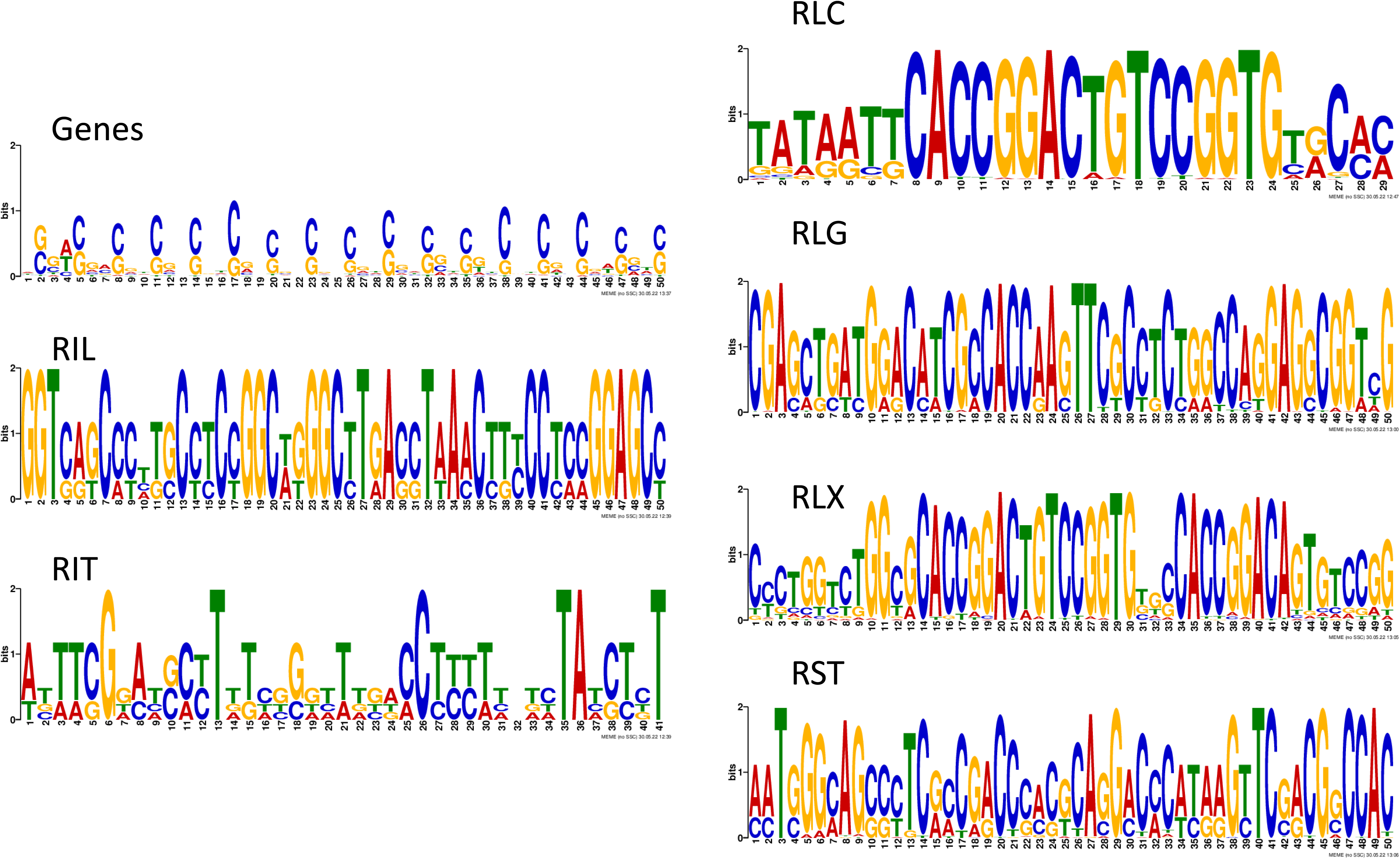
Overrepresented motifs in structured/low MFE regions (<-40 kcal/mol) of structured features. Structured regions of each superfamily were entered into MEME motif finder (See Methods), and logos represent the most highly overrespresented motif found in each superfamily.

**Figure S6.**
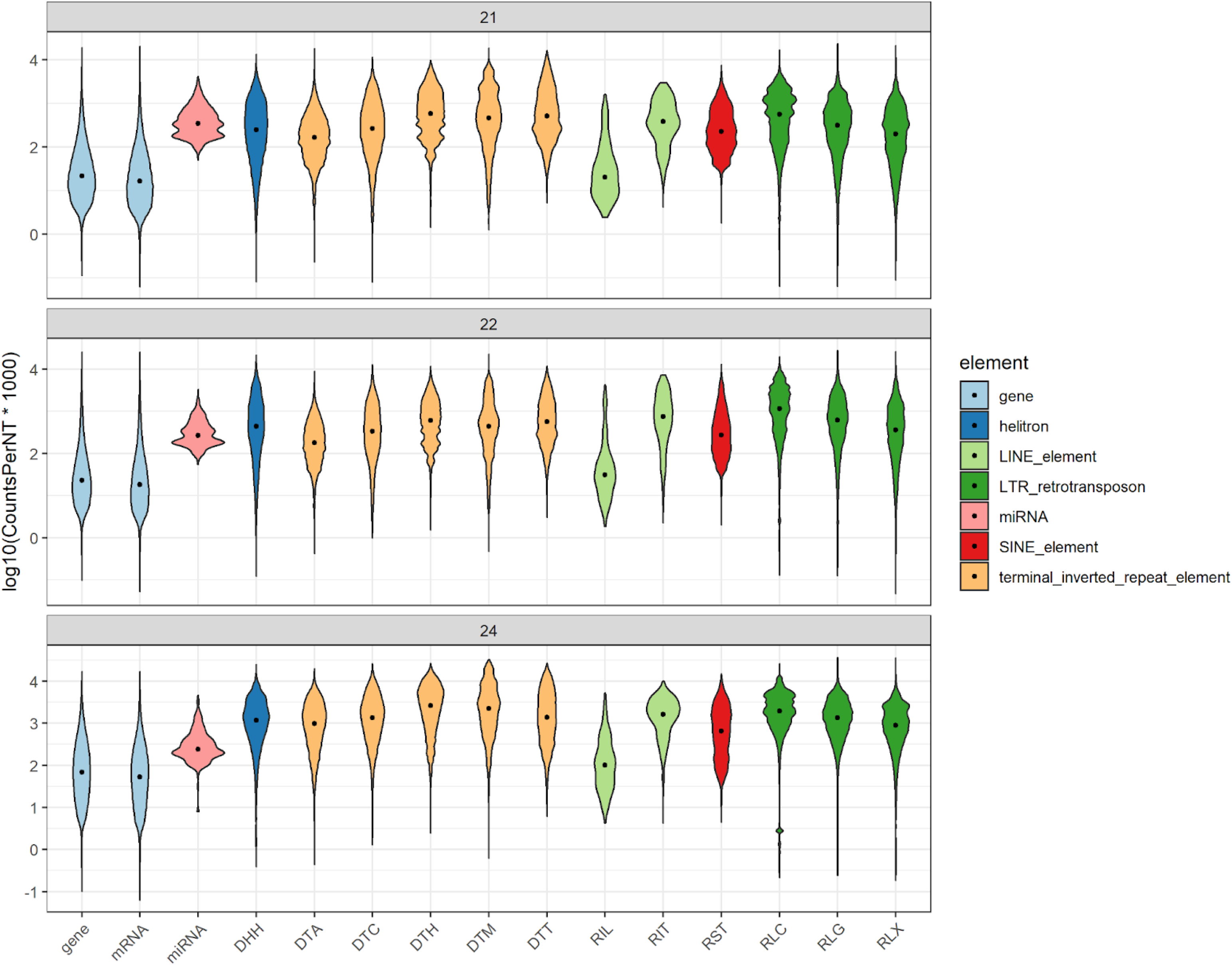
Variation in siRNA mapping between feature types. Violin plots show the distributions of siRNA mapping densities in log10(siRNA species counts per kilobase) for each superfamily/genomic feature, and black dots show the mean of the distribution. Panels represent siRNA size classes (21-nt, 22-nt, 24-nt).

**Figure S7.**
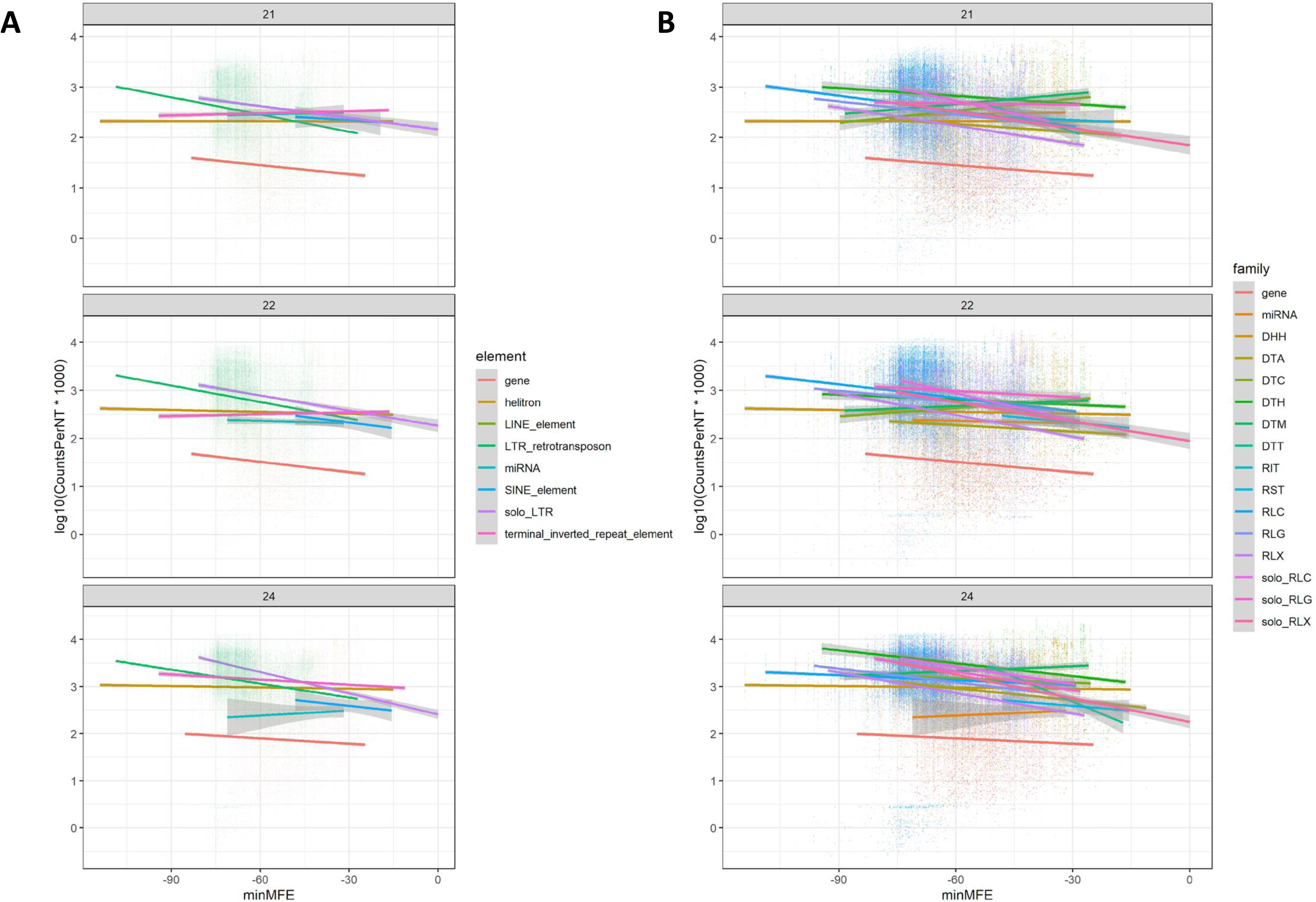
Linear models of minMFE vs siRNA mapping density. Regressions show siRNA species mapping density in log10(siRNA species counts per kb) as a function of minMFE. The left column describes these relationships in broader categories (LTRs, TIRs, etc), while the right column shows individual superfamilies. Effectively, these graphs depict the same information found in Table S2. P-values and R2 values can be found in Table S2.

**Figure S8.**
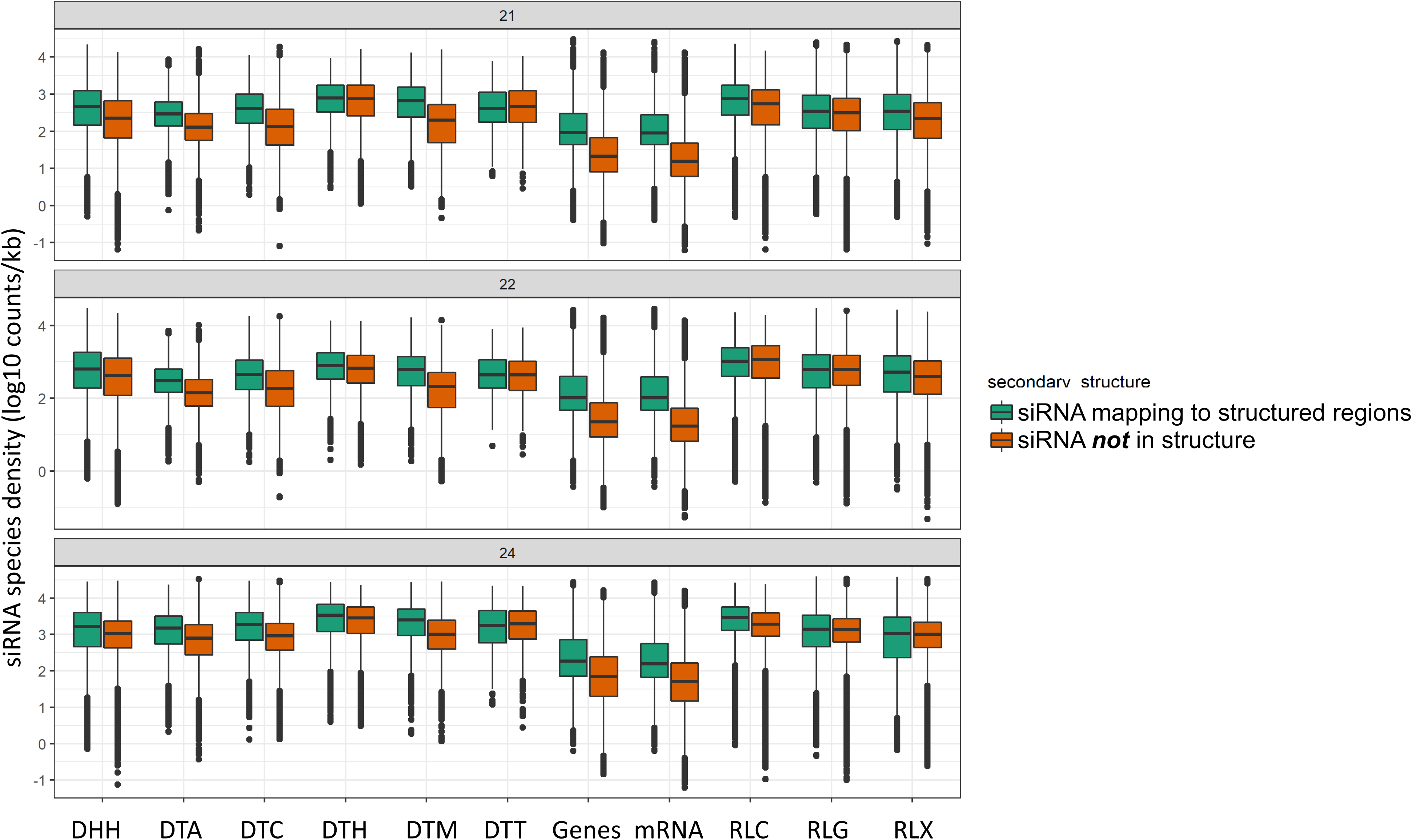
siRNA species mapping density in structured regions vs unstructured regions. For each structured feature (minMFE significantly lower than mean randomized minMFE (see **Methods**), siRNAs mapping to the feature were divided into those mapping to structured regions (<-40 kcal/mol) and those outside of structured regions. Boxplot central lines show the median, and boxes show the 25% and 75% quartiles. Statistical significance in these comparisons can be seen in Table S3.

**Figure S9.**
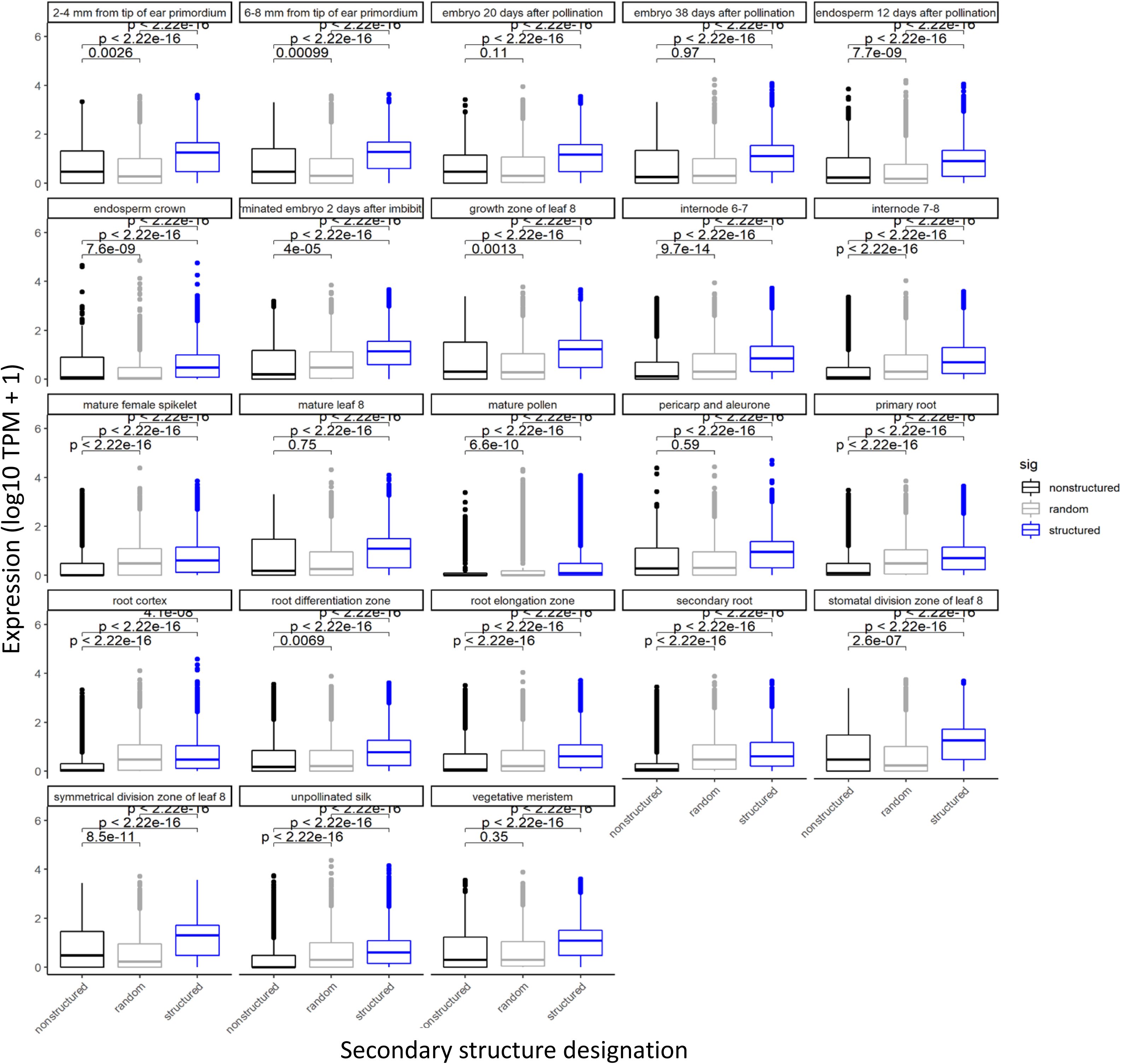
Expression between structured, random, and unstructured genes in 23 B73 tissues. Expression is represented in log10 TPM+1, and structure designations are from the primary sequences in B73 (see **Methods**). Expression data are from Walley et al., 2016 and were downloaded from the ATLAS expression database (E-GEOD-50191). Boxplot central lines represent the median, and boxes represent the 25% and 75% quartiles.

**Figure S10.**
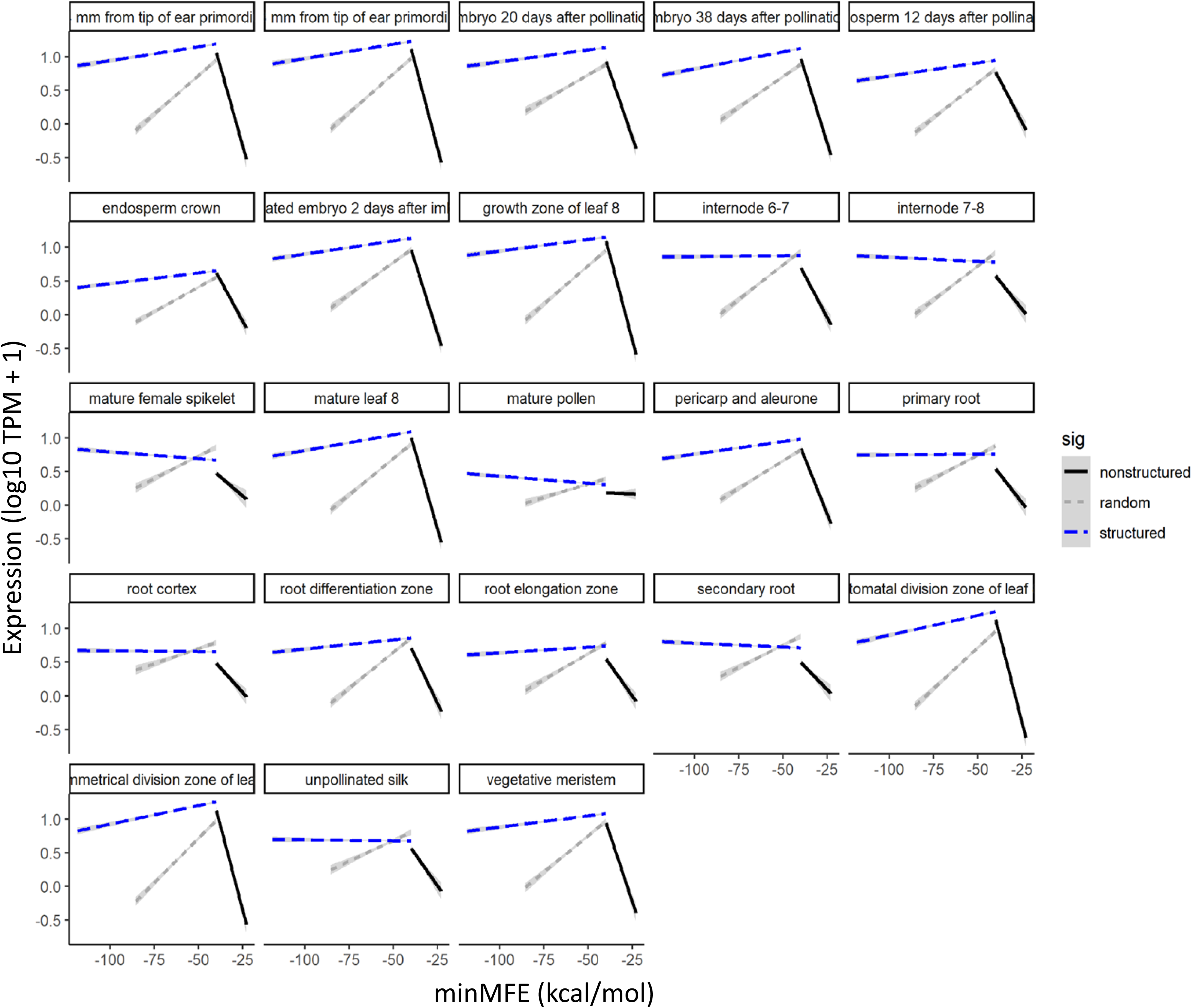
Expression as a function of minMFE in 23 B73 tissues. Expression is represented in log10 TPM+1, and structure designations are from the primary sequences in B73 (see **Methods**). Expression data are from Walley et al., 2016 and were downloaded from the ATLAS expression database (E-GEOD-50191).

**Figure S11.**
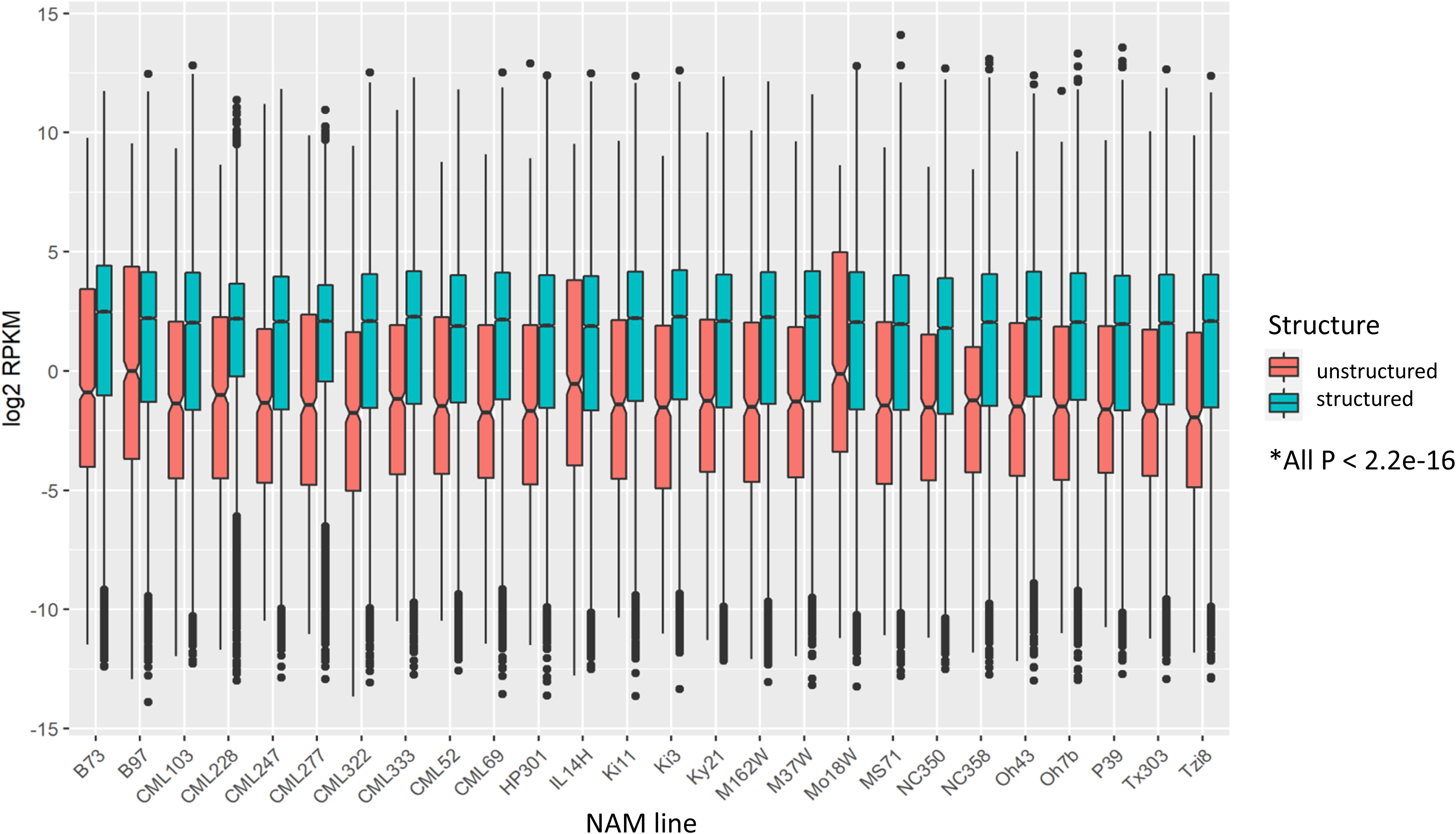
Expression of structured vs unstructured genes in the 26 NAM lines. Expression is represented in log2 RPKM and is from Hufford et al. 2021 Structure designations were inferred from the primary sequences in B73 (see **Methods**).

**Table S1.** Small RNA libraries used

| **Library** | **Tissue** | **Paper** |
| --- | --- | --- |
| SRR3684389 | Kernel | Huanhuan H et al., |
| SRR3684242 | Kernel | Huanhuan H et al., |
| SRR2086100 | Leaf | Lunardon A et al., 2014 |
| SRR2086104 | Leaf | Lunardon A et al., 2014 |
| SRR2086116 | Leaf | Lunardon A et al., 2014 |
| SRR2086120 | Leaf | Lunardon A et al., 2014 |
| SRR2086132 | Leaf | Lunardon A et al., 2014 |
| SRR2086136 | Leaf | Lunardon A et al., 2014 |
| SRR2106180 | Leaf | Lunardon A et al., 2014 |
| SRR2106184 | Leaf | Lunardon A et al., 2014 |
| SRR2106196 | Leaf | Lunardon A et al., 2014 |
| SRR2106200 | Leaf | Lunardon A et al., 2014 |
| SRR1917157 | Imbibed kernels | Petsch K et al., 2015 |
| SRR1917158 | Imbibed kernels | Petsch K et al., 2015 |
| SRR895785 | Ears (female inflorescences) | Liu H et al., 2014 |
| SRR1186264 | Leaf; 3rd and 4th | Diez et al., 2014 |
| SRR032091 | Tassels | Zhang L et al., 2009 |
| SRR032090 | Seedling | Zhang L et al., 2009 |
| SRR032089 | Roots | Zhang L et al., 2009 |
| SRR032088 | Pollen | Zhang L et al., 2009 |
| SRR032087 | Ear | Zhang L et al., 2009 |
| GSM306487 | Immature ear | Arteaga-Vazques et al., 2010 |
| GSM306488 | Immature ear | Arteaga-Vazques et al., 2010 |
| SRX483603 | Leaf | Diez et al., 2014 |

**Table S2.** Correlations between metrics of predicted secondary structure and siRNA mapping density within feature types

| Feature | Structure measurement | 21-nt siRNA | 22-nt siRNA | 24-nt siRNA |
| --- | --- | --- | --- | --- |
| DHH | minMFE | P = 3.070e-78 R2 = 7.100e-03 | P = 2.180e-117 R2 = 1.070e-02 | P = 3.950e-103 R2 = 9.400e-03 |
|  | meanMFE | P = 0.000e+00 R2 = 7.970e-02 | P = 0.000e+00 R2 = 6.990e-02 | P = 0.000e+00 R2 = 9.520e-02 |
|  | MFEvariance | P = 1.260e-43 R2 = 3.890e-03 | P = 5.670e-25 R2 = 2.160e-03 | P = 2.380e-169 R2 = 1.550e-02 |
|  | PctMFEltthresh | P = 6.370e-226 R2 = 2.070e-02 | P = 9.220e-134 R2 = 1.220e-02 | P = 0.000e+00 R2 = 3.910e-02 |
| DTA | minMFE | P = 4.770e-198 R2 = 1.490e-01 | P = 8.580e-194 R2 = 1.460e-01 | P = 3.610e-109 R2 = 8.420e-02 |
|  | meanMFE | P = 1.610e-150 R2 = 1.150e-01 | P = 4.770e-133 R2 = 1.020e-01 | P = 2.280e-48 R2 = 3.740e-02 |
|  | MFEvariance | P = 5.410e-148 R2 = 1.130e-01 | P = 3.840e-151 R2 = 1.150e-01 | P = 3.630e-130 R2 = 9.990e-02 |
|  | PctMFEltthresh | P = 1.350e-185 R2 = 1.400e-01 | P = 1.030e-175 R2 = 1.330e-01 | P = 4.910e-75 R2 = 5.820e-02 |
| DTC | minMFE | P = 1.360e-12 R2 = 3.900e-02 | P = 9.750e-18 R2 = 5.660e-02 | P = 4.440e-28 R2 = 9.120e-02 |
|  | meanMFE | P = 6.710e-68 R2 = 2.140e-01 | P = 2.940e-48 R2 = 1.550e-01 | P = 1.920e-41 R2 = 1.340e-01 |
|  | MFEvariance | P = 1.520e-03 R2 = 7.940e-03 | P = 7.770e-07 R2 = 1.920e-02 | P = 2.490e-12 R2 = 3.810e-02 |
|  | PctMFEltthresh | P = 1.890e-76 R2 = 2.380e-01 | P = 9.480e-56 R2 = 1.780e-01 | P = 1.120e-46 R2 = 1.510e-01 |
| DTH | minMFE | P = 2.990e-76 R2 = 6.640e-02 | P = 2.740e-48 R2 = 4.200e-02 | P = 1.460e-32 R2 = 2.800e-02 |
|  | meanMFE | P = 1.800e-69 R2 = 6.060e-02 | P = 5.940e-33 R2 = 2.840e-02 | P = 2.800e-13 R2 = 1.070e-02 |
|  | MFEvariance | P = 7.420e-04 R2 = 2.290e-03 | P = 2.360e-07 R2 = 5.360e-03 | P = 2.520e-14 R2 = 1.160e-02 |
|  | PctMFEltthresh | P = 1.630e-38 R2 = 3.330e-02 | P = 2.050e-21 R2 = 1.800e-02 | P = 5.810e-08 R2 = 5.910e-03 |
| DTM | minMFE | P = 6.100e-04 R2 = 8.880e-03 | **P = 9.640e-01** R2 = 1.550e-06 | P = 4.210e-04 R2 = 9.400e-03 |
|  | meanMFE | P = 1.220e-68 R2 = 2.080e-01 | P = 1.830e-76 R2 = 2.290e-01 | P = 2.080e-48 R2 = 1.500e-01 |
|  | MFEvariance | **P = 2.690e-01** R2 = 9.260e-04 | **P = 6.550e-01** R2 = 1.510e-04 | P = 1.490e-03 R2 = 7.630e-03 |
|  | PctMFEltthresh | P = 2.980e-10 R2 = 2.970e-02 | P = 2.650e-11 R2 = 3.320e-02 | P = 1.720e-02 R2 = 4.300e-03 |
| DTT | minMFE | P = 1.480e-12 R2 = 1.040e-01 | P = 3.250e-12 R2 = 1.010e-01 | P = 9.380e-21 R2 = 1.740e-01 |
|  | meanMFE | P = 2.260e-05 R2 = 3.870e-02 | P = 3.680e-04 R2 = 2.750e-02 | P = 1.750e-09 R2 = 7.650e-02 |
|  | MFEvariance | P = 3.040e-05 R2 = 3.750e-02 | P = 7.770e-06 R2 = 4.290e-02 | P = 1.630e-08 R2 = 6.760e-02 |
|  | PctMFEltthresh | **P = 2.200e-01** R2 = 3.290e-03 | **P = 5.340e-01** R2 = 8.510e-04 | P = 1.500e-03 R2 = 2.190e-02 |
| gene | minMFE | P = 5.560e-129 R2 = 1.480e-02 | P = 4.120e-106 R2 = 1.210e-02 | **P = 7.050e-02** R2 = 8.350e-05 |
|  | meanMFE | P = 2.850e-124 R2 = 1.420e-02 | P = 2.520e-158 R2 = 1.820e-02 | P = 0.000e+00 R2 = 6.820e-02 |
|  | MFEvariance | P = 8.180e-71 R2 = 8.050e-03 | P = 5.990e-48 R2 = 5.390e-03 | P = 7.650e-06 R2 = 5.110e-04 |
|  | PctMFEltthresh | P = 3.100e-144 R2 = 1.660e-02 | P = 3.590e-185 R2 = 2.130e-02 | P = 0.000e+00 R2 = 6.910e-02 |
| miRNA | minMFE | **P = 2.290e-01** R2 = 1.380e-02 | **P = 9.360e-02** R2 = 2.650e-02 | **P = 4.180e-01** R2 = 6.270e-03 |
| | meanMFE | $P = 3.190e-01$ $R^2 = 9.470e-03$ | $P = 8.550e-02$ $R^2 = 2.790e-02$ | $P = 1.360e-01$ $R^2 = 2.100e-02$ |
| | MFEvariance | $P = 1.640e-01$ $R^2 = 1.840e-02$ | $P = 4.420e-01$ $R^2 = 5.640e-03$ | $P = 2.060e-01$ $R^2 = 1.520e-02$ |
| | PctMFEltthresh | $P = 1.340e-01$ $R^2 = 2.130e-02$ | $P = 5.050e-02$ $R^2 = 3.590e-02$ | $P = 7.070e-02$ $R^2 = 3.080e-02$ |
| RIL | minMFE | $P = 4.860e-02$ $R^2 = 1.100e-01$ | **$P = 6.800e-02$** $R^2 = 9.460e-02$ | $P = 1.240e-03$ $R^2 = 2.670e-01$ |
| | meanMFE | $P = 2.780e-02$ $R^2 = 1.340e-01$ | $P = 2.960e-02$ $R^2 = 1.320e-01$ | $P = 3.180e-04$ $R^2 = 3.210e-01$ |
| | MFEvariance | **$P = 7.340e-02$** $R^2 = 9.120e-02$ | **$P = 9.180e-02$** $R^2 = 8.130e-02$ | **$P = 5.010e-02$** $R^2 = 1.080e-01$ |
| | PctMFEltthresh | **$P = 6.620e-01$** $R^2 = 5.670e-03$ | **$P = 6.090e-01$** $R^2 = 7.760e-03$ | **$P = 3.120e-01$** $R^2 = 3.000e-02$ |
| RIT | minMFE | **$P = 3.090e-01$** $R^2 = 3.820e-02$ | **$P = 2.380e-01$** $R^2 = 5.120e-02$ | **$P = 4.010e-01$** $R^2 = 2.620e-02$ |
| | meanMFE | **$P = 1.260e-01$** $R^2 = 8.460e-02$ | **$P = 8.900e-02$** $R^2 = 1.030e-01$ | **$P = 2.660e-01$** $R^2 = 4.560e-02$ |
| | MFEvariance | $P = 2.300e-03$ $R^2 = 2.960e-01$ | $P = 2.410e-03$ $R^2 = 2.930e-01$ | $P = 2.750e-02$ $R^2 = 1.670e-01$ |
| | PctMFEltthresh | **$P = 2.260e-01$** $R^2 = 5.380e-02$ | **$P = 1.470e-01$** $R^2 = 7.630e-02$ | **$P = 2.720e-01$** $R^2 = 4.450e-02$ |
| RLC | minMFE | $P = 0.000e+00$ $R^2 = 6.980e-02$ | $P = 0.000e+00$ $R^2 = 6.990e-02$ | $P = 0.000e+00$ $R^2 = 1.140e-01$ |
| | meanMFE | $P = 0.000e+00$ $R^2 = 7.130e-02$ | $P = 0.000e+00$ $R^2 = 6.270e-02$ | $P = 0.000e+00$ $R^2 = 9.500e-02$ |
| | MFEvariance | $P = 0.000e+00$ $R^2 = 6.890e-02$ | $P = 0.000e+00$ $R^2 = 5.660e-02$ | $P = 0.000e+00$ $R^2 = 1.310e-01$ |
| | PctMFEltthresh | $P = 3.700e-172$ $R^2 = 1.750e-02$ | $P = 4.310e-110$ $R^2 = 1.120e-02$ | $P = 3.530e-214$ $R^2 = 2.180e-02$ |
| RLG | minMFE | $P = 0.000e+00$ $R^2 = 7.040e-02$ | $P = 0.000e+00$ $R^2 = 5.370e-02$ | $P = 0.000e+00$ $R^2 = 1.550e-01$ |
| | meanMFE | $P = 0.000e+00$ $R^2 = 1.600e-01$ | $P = 0.000e+00$ $R^2 = 1.080e-01$ | $P = 0.000e+00$ $R^2 = 2.410e-01$ |
| | MFEvariance | $P = 1.350e-223$ $R^2 = 1.440e-02$ | $P = 7.300e-160$ $R^2 = 1.030e-02$ | $P = 0.000e+00$ $R^2 = 7.240e-02$ |
| | PctMFEltthresh | $P = 0.000e+00$ $R^2 = 1.360e-01$ | $P = 0.000e+00$ $R^2 = 8.430e-02$ | $P = 0.000e+00$ $R^2 = 2.500e-01$ |
| RLX | minMFE | $P = 0.000e+00$ $R^2 = 1.100e-01$ | $P = 0.000e+00$ $R^2 = 1.280e-01$ | $P = 0.000e+00$ $R^2 = 1.480e-01$ |
|  | meanMFE | P = 1.050e-275<br>R2 = 7.480e-02 | P = 1.690e-254<br>R2 = 6.920e-02 | P = 0.000e+00<br>R2 = 9.460e-02 |
|  | MFEvariance | P = 0.000e+00<br>R2 = 9.380e-02 | P = 9.280e-319<br>R2 = 8.600e-02 | P = 0.000e+00<br>R2 = 1.290e-01 |
|  | PctMFEltthresh | P = 1.970e-133<br>R2 = 3.660e-02 | P = 5.920e-106<br>R2 = 2.910e-02 | P = 5.420e-211<br>R2 = 5.760e-02 |
| RST | minMFE | P = 2.240e-41 R2 = 1.750e-01 | P = 4.840e-30<br>R2 = 1.280e-01 | P = 2.780e-46<br>R2 = 1.940e-01 |
|  | meanMFE | P = 7.040e-45 R2 = 1.890e-01 | P = 1.280e-30<br>R2 = 1.310e-01 | P = 8.970e-51<br>R2 = 2.120e-01 |
|  | MFEvariance | P = 6.170e-08 R2 = 3.060e-02 | P = 3.770e-05<br>R2 = 1.780e-02 | P = 2.910e-07<br>R2 = 2.750e-02 |
|  | PctMFEltthresh | P = 2.970e-22 R2 = 9.490e-02 | P = 1.220e-12<br>R2 = 5.210e-02 | P = 3.580e-29<br>R2 = 1.250e-01 |
| solo_RLC | minMFE | P = 5.310e-60 R2 = 2.950e-01 | P = 1.010e-51<br>R2 = 2.590e-01 | P = 2.970e-57<br>R2 = 2.830e-01 |
|  | meanMFE | P = 6.590e-57 R2 = 2.810e-01 | P = 3.310e-24<br>R2 = 1.260e-01 | P = 3.480e-33<br>R2 = 1.720e-01 |
|  | MFEvariance | P = 3.330e-23 R2 = 1.210e-01 | P = 1.310e-19<br>R2 = 1.020e-01 | P = 8.520e-27<br>R2 = 1.390e-01 |
|  | PctMFEltthresh | P = 2.110e-47 R2 = 2.390e-01 | P = 1.170e-16<br>R2 = 8.590e-02 | P = 1.720e-25<br>R2 = 1.330e-01 |
| solo_RLG | minMFE | P = 6.900e-54 R2 = 8.150e-02 | P = 1.710e-65<br>R2 = 9.880e-02 | P = 8.620e-178<br>R2 = 2.500e-01 |
|  | meanMFE | P = 2.620e-117<br>R2 = 1.720e-01 | P = 6.950e-101<br>R2 = 1.490e-01 | P = 1.460e-190<br>R2 = 2.660e-01 |
|  | MFEvariance | P = 1.680e-09 R2 = 1.280e-02 | P = 6.860e-06<br>R2 = 7.180e-03 | P = 8.990e-02<br>R2 = 1.020e-03 |
|  | PctMFEltthresh | P = 1.480e-84 R2 = 1.260e-01 | P = 1.690e-70<br>R2 = 1.060e-01 | P = 3.420e-163<br>R2 = 2.320e-01 |
| solo_RLX | minMFE | P = 3.290e-99 R2 = 1.800e-01 | P = 4.640e-105<br>R2 = 1.900e-01 | P = 1.140e-159<br>R2 = 2.750e-01 |
|  | meanMFE | P = 2.230e-82 R2 = 1.520e-01 | P = 2.100e-85<br>R2 = 1.570e-01 | P = 2.590e-120<br>R2 = 2.150e-01 |
|  | MFEvariance | P = 6.470e-44 R2 = 8.230e-02 | P = 3.240e-26<br>R2 = 4.870e-02 | P = 2.760e-67<br>R2 = 1.250e-01 |
|  | PctMFEltthresh | P = 2.920e-50 R2 = 9.410e-02 | P = 1.610e-50<br>R2 = 9.460e-02 | P = 6.050e-77<br>R2 = 1.420e-01 |
| All features | minMFE | P = 0.000e+00<br>R2 = 9.060e-02 | P = 0.000e+00<br>R2 = 1.030e-01 | P = 0.000e+00<br>R2 = 7.380e-02 |
|  | meanMFE | P = 0.000e+00<br>R2 = 1.660e-02 | P = 0.000e+00<br>R2 = 8.610e-03 | P = 5.010e-227<br>R2 = 4.310e-03 |
|  | MFEvariance | P = 0.000e+00<br>R2 = 1.790e-02 | P = 0.000e+00<br>R2 = 1.700e-02 | P = 0.000e+00<br>R2 = 2.350e-02 |
|  | PctMFEltthresh | P = 2.950e-10 R2 = 1.660e-04 | P = 2.130e-47<br>R2 = 8.730e-04 | P = 2.410e-52<br>R2 = 9.670e-04 |

**Table S3.** Statistics from mixed-effect models comparing siRNA species mapping density between structured and unstructured regions (i.e., regions less than and greater than −40 kcal/mol) of structured features (i.e., features with significantly lower minMFE than five randomizations; see **Methods**)

| Feature | 21-nt siRNA | 22-nt siRNA | 24-nt siRNA |
| --- | --- | --- | --- |
| RLC | Estimate = -2.400e-01 \|<br>P = 0.000e+00 \| std. err.<br>= 1.860e-03 | Estimate = 1.860e-01<br>\| P = 0.000e+00 \|<br>std. err. = 1.750e-03 | Estimate = -2.190e-01 \| P = 0.000e+00<br>\| std. err. = 1.320e-03 |
| RLG | Estimate = 1.580e-01 \| P<br>= 0.000e+00 \| std. err. =<br>1.650e-03 | Estimate = 3.000e-01<br>\| P = 0.000e+00 \|<br>std. err. = 1.520e-03 | Estimate = 2.070e-01 \| P = 0.000e+00<br>\| std. err. = 1.230e-03 |
| RLX | Estimate = -1.780e-01 \|<br>P = 0.000e+00 \| std. err.<br>= 4.140e-03 | Estimate = 5.720e-02<br>\| P = 5.350e-47 \| std.<br>err. = 3.970e-03 | Estimate = 2.670e-01 \| P = 0.000e+00<br>\| std. err. = 3.140e-03 |
| DHH | Estimate = -2.830e-01 \|<br>P = 0.000e+00 \| std. err.<br>= 2.280e-03 | Estimate = 1.230e-02<br>\| P = 1.930e-08 \| std.<br>err. = 2.190e-03 | Estimate = -7.890e-02 \| P = 0.000e+00<br>\| std. err. = 1.680e-03 |
| DTC | Estimate = -5.640e-01 \|<br>P = 9.980e-279 \| std.<br>err. = 1.550e-02 | Estimate = -3.370e-01 \| P = 7.810e-101 \|<br>std. err. = 1.570e-02 | Estimate = -3.630e-01 \| P = 8.310e-208 \| std. err. = 1.170e-02 |
| DTA | Estimate = -6.380e-01 \|<br>P = 0.000e+00 \| std. err.<br>= 8.910e-03 | Estimate = -5.760e-01 \| P = 0.000e+00 \|<br>std. err. = 8.500e-03 | Estimate = -4.450e-01 \| P = 0.000e+00<br>\| std. err. = 6.610e-03 |
| DTH | Estimate = -1.270e-01 \|<br>P = 6.240e-28 \| std. err.<br>= 1.160e-02 | Estimate = -2.100e-01 \| P = 1.610e-75 \|<br>std. err. = 1.140e-02 | Estimate = -2.630e-01 \| P = 5.490e-209 \| std. err. = 8.470e-03 |
| DTM | Estimate = -9.740e-01 \|<br>P = 0.000e+00 \| std. err.<br>= 1.630e-02 | Estimate = -8.830e-01 \| P = 0.000e+00 \|<br>std. err. = 1.520e-02 | Estimate = -7.690e-01 \| P = 0.000e+00<br>\| std. err. = 1.230e-02 |
| DTT | Estimate = -3.180e-01 \|<br>P = 7.850e-15 \| std. err.<br>= 4.060e-02 | Estimate = -4.600e-01 \| P = 6.030e-31 \|<br>std. err. = 3.920e-02 | Estimate = -4.430e-01 \| P = 8.540e-50<br>\| std. err. = 2.940e-02 |
| gene | Estimate = -8.780e-01 \|<br>P = 0.000e+00 \| std. err.<br>= 5.980e-03 | Estimate = -7.910e-01 \| P = 0.000e+00 \|<br>std. err. = 5.800e-03 | Estimate = -5.400e-01 \| P = 0.000e+00<br>\| std. err. = 4.340e-03 |
| mRNA | Estimate = -1.080e+00 \|<br>P = 0.000e+00 \| std. err.<br>= 3.830e-03 | Estimate = -<br>1.020e+00 \| P =<br>0.000e+00 \| std. err.<br>= 3.720e-03 | Estimate = -6.840e-01 \| P = 0.000e+00<br>\| std. err. = 2.870e-03 |

**Table S4.** Illumina adapter sequences used to trim each small RNA library with CutAdapt (see **Methods**)

| Library | Adapter |
| --- | --- |
| GSM306487 | CTGTAGG |
| GSM306488 | CTGTAGG |
| SRR032087 | CTGTAGG |
| SRR032088 | CTGTAGG |
| SRR032089 | CTGTAGG |
| SRR032090 | CTGTAGG |
| SRR032091 | CTGTAGG |
| SRR1186264 | ATCTCGT |
| SRR1917157 | AGATCGG |
| SRR1917158 | AGATCGG |
| SRR2086100 | TGGAATT |
| SRR2086104 | TGGAATT |
| SRR2086116 | TGGAATT |
| SRR2086120 | TGGAATT |
| SRR2086132 | TGGAATT |
| SRR2086136 | TGGAATT |
| SRR2106180 | TGCAGCA |
| SRR2106184 | TGCAGCA |
| SRR2106196 | TGCAGCA |
| SRR2106200 | TGCAGCA |
| SRR3684242 | TCGTATG |
| SRR3684389 | TCGTATG |
| SRR895785 | TCGTATG |
| SRX483603 | ATCTCGT |

